# CTCF sites display cell cycle dependent dynamics in factor binding and nucleosome positioning

**DOI:** 10.1101/365866

**Authors:** Marlies E. Oomen, Anders S. Hansen, Yu Liu, Xavier Darzacq, Job Dekker

**Author notes:** Correspondence to: Job Dekker.

## Abstract

CTCF plays a key role in formation of topologically associating domains (TADs) and loops in interphase. During mitosis TADs are absent, but how TAD formation is dynamically controlled during the cell cycle is not known. Several contradicting observations have been made regarding CTCF binding to mitotic chromatin using both genomics and microscopy-based techniques. Here we have used 4 different assays to address this debate. First, using 5C we confirmed that TADs and CTCF loops are readily detected in interphase, but absent during prometaphase. Second, ATAC-seq analysis showed that CTCF sites display greatly reduced accessibility and lose the CTCF footprint in prometaphase, suggesting loss of CTCF binding and rearrangement of the nucleosomal array around the binding motif. In contrast, transcription start sites remain accessible in prometaphase, although adjacent nucleosomes can also become repositioned and occupy at least a subset of start sites during mitosis. Third, loss of site-specific CTCF binding was directly demonstrated using CUT&RUN. Histone modifications and histone variants are maintained in mitosis, suggesting a role in bookmarking of active CTCF sites. Finally, live-cell imaging, fluorescence recovery after photobleaching and single molecule tracking showed that almost all CTCF chromatin binding is lost in prometaphase. Combined, our results demonstrate loss of CTCF binding to CTCF sites during prometaphase and rearrangement of the chromatin landscape around CTCF motifs. This contributes to loss of TADs and CTCF loops during mitosis, and reveals that CTCF sites, a key architectural cis-element of the genome, display cell cycle stage-dependent dynamics in factor binding and nucleosome positioning.

## Introduction

Several studies have observed a key role of CCCTC-binding factor (CTCF) in organizing the linear genome in topologically associating domains (TADs) and loops in interphase vertebrate cells (Dixon et al. 2012; Nora et al. 2017, 2012). CTCF is an 11 zinc finger protein that binds a well-defined motif to which it can bind only in one direction (Kim et al. 2007). Nucleosomes flanking CTCF bound sites are strongly positioned (Fu et al. 2008). In addition, flanking nucleosomes contain histone modifications such as H3K4 methylation and histone variants such as H2A.Z (Jin et al. 2009; Nekrasov et al. 2012). Although CTCF has about 42,000 predicted binding sites in the human genome, only a subset of CTCF sites are bound in a given cell type.

It has been proposed that topologically associating domains (TADs) and CTCF-loops are formed as a result of cohesin-dependent loop extrusion (Dekker and Mirny 2016; Fudenberg et al. 2018; Hansen et al. 2017). According to this model, when cohesin is loaded on the chromatin, it will be able to form a loop between two loci and will keep extruding until it is blocked by CTCF, which will function as a boundary element. Whether cohesin is blocked by bound CTCF and whether two CTCF-occupied sites can form a loop depends on the orientation of the CTCF motif: looping is mostly observed between CTCF sites in a convergent orientation (Vietri Rudan et al. 2015; Wit et al. 2015; Rao et al. 2014; Guo et al. 2015). CTCF loops often define TADs that are implicated in gene regulation. In one model, cohesin-dependent loop extrusion can bring together enhancers and promoters located in the same TAD, but elements located in different TADs cannot be similarly juxtaposed due to the blocking activity of CTCF-bound sites (Phillips-Cremins and Corces 2013; Dekker and Mirny 2016).

Chromosome organization changes dramatically during mitosis (Flemming 1878; Gibcus et al. 2018; Naumova et al. 2013; Nagano et al. 2017; Earnshaw and Laemmli 1983; Marsden and Laemmli 1979). The structural features of interphase chromosomes described by 5C and Hi-C, such as TADs and A- and B-compartments are lost in prometaphase (Naumova et al. 2013). Current models, based on modeling Hi-C data for prometaphase cells combined with extensive earlier imaging data (Marsden and Laemmli 1979; Earnshaw and Laemmli 1983; Adolph 1980), propose that mitotic chromosomes are organized as arrays of nested loops that are helically arranged around a spiraling central axis (Gibcus et al. 2018). These loops can be generated by a process of loop extrusion mediated by condensin complexes (Gibcus et al. 2018; Dekker and Mirny 2016; Fudenberg et al. 2018; Goloborodko et al. 2016).

Although it is clear that loss of CTCF causes genome-wide loss of TADs in interphase (Nora et al. 2017), whether the loss of TADs during mitosis is due to regulation of CTCF is currently unclear. First, it is possible that condensin-mediated loop extrusion, unlike cohesin-mediated extrusion, is not blocked by CTCF. Second, it is possible that even when condensin is blocked by CTCF, formation of packed arrays of loops in mitosis makes detection of such boundaries difficult. Indeed, polymer simulations have shown that extensive condensin-mediated loop extrusion could lead to loss of TADs as detected by Hi-C, even when CTCF is still bound and blocks extrusion (Gibcus et al. 2018). Alternatively, TAD boundaries could be absent because CTCF dissociates from chromatin during mitosis. Along these lines, CTCF becomes highly phosphorylated in mitosis (Dephoure et al. 2008; Dovat et al. 2002; Rizkallah and Hurt 2009) and in vitro assays show that DNA binding capability of phosphorylated CTCF is dramatically reduced (Sekiya et al. 2017; Jantz and Berg 2004).

There have been several studies to examine chromatin/protein factor binding in mitotic cells over the past decades using both microscopy and genomics techniques such as ChIP-seq and DNase I sensitivity assays. Several studies suggest that most factors lose site-specific binding to the chromatin during mitosis (Hsiung et al. 2015b; Martinez-Balbas et al. 1995). However, other studies, mainly using imaging or western blot analysis of chromosome associated proteins, report maintenance of factor binding in mitosis (Teves et al. 2016; Burke et al. 2005; Liu et al. 2017; Chen et al. 2005). There are several possible reasons that could explain why these studies show conflicting results. First, it has been shown that formaldehyde fixation can affect protein association with mitotic chromosomes, and therefore prevent observation of factor binding by both microscopy and ChIP-seq (Teves et al. 2016; Pallier et al. 2003). Additionally, when performing population wide genomic studies in mitosis, cells need to be synchronized using drugs, cell sorting or modified cell lines (Hochegger et al. 2007; Gibcus et al. 2018; Taylor 2004; Jackman and O’Connor 2001). It is very important to obtain pure synchronized populations, as contamination of interphase cells, especially in studies using immunoprecipitation, can lead to an overestimation of signal in mitosis. Furthermore, although microscopy definitely has advantages over population wide studies, microscopy does not capture information on site specific binding of factors. Moreover, it is important to distinguish co-localization of factors with the mitotic chromatin from site specific binding, which could function as a mitotic bookmark (Raccaud and Suter 2018). Although it remains unresolved whether specific factors maintain or lose binding throughout the cell cycle, it is generally accepted that mitotic chromatin partially maintains regions of hyper-accessibility (Teves et al. 2016; Hsiung et al. 2015a; Martinez-Balbas et al. 1995; Teves et al. 2018).

In this study, we used a combination of several genomics techniques and live cell imaging to study cell cycle dynamics of CTCF mediated looping interactions, CTCF binding and local chromatin state flanking CTCF binding sites. We find that overall CTCF binding is temporarily lost during prometaphase. Loss of CTCF binding could contribute to the loss of TADs in mitosis. We also find that upon CTCF loss nucleosomes lose their strong positioning with respect to the CTCF sites, and rearrange while maintaining a regular spacing and start to occupy the CTCF motif. While transcription start sites (TSS) remain generally accessible, changes in nucleosome spacing and positioning can also occur in these regions during mitosis. Finally, specific histone variants and histone modifications that are present at CTCF sites and TSSs in interphase, such as H2A.Z and H3K4 methylation are maintained throughout the cell cycle. Taken together, these findings reveal extensive chromatin dynamics at CTCF sites during the cell cycle that correlate with TAD and loop formation and dissolution, and suggest roles for histone modifications and variants in mitotic bookmarking at these CTCF binding sites.

## Results

### 5C shows loss of TADs and CTCF loops in prometaphase

It has been shown that interphase structures detected by 3C-based methods, such as TADs and compartments, are lost in mitosis (Nagano et al. 2017; Gibcus et al. 2018; Naumova et al. 2013; Oomen and Dekker 2017). However, the resolution of these previous studies was not sufficient to investigate specific looping interactions, e.g. between CTCF sites. We therefore applied a targeted 5C approach (Dostie et al. 2006) that allows high resolution analysis (10 to 15 kb) for domains up to several megabases in interphase and mitosis. We chose to use a validated 5C primer set that targets two 2 MB regions that we previously analyzed and where we identified several TADs and CTCF loops in non-synchronous cell cultures (Hnisz et al. 2016). We synchronized HeLa S3 cells by first arresting cells in early S-phase using a thymidine block, followed by an arrest in prometaphase using nocodazole (Naumova et al. 2013). We confirmed the cell cycle state of the non-synchronous and mitotic (prometaphase) cell populations using flow cytometry (Supplemental Fig. S1) and quantified the mitotic index (percentage of cells with condensed chromosomes) using fluorescence microscopy of DAPI stained cells. HeLa S3 asynchronous populations typically had a mitotic index count of approximately 5%, whereas a nocodazole arrested culture contained 95-98% mitotic cells. In addition to using mitotic cells, we also biochemically purified mitotic chromatin from prometaphase cells (Gasser and Laemmli 1987). Using a percoll gradient, we separated mitotic chromosomes from interphase nuclei, resulting in a pure mitotic chromatin preparation. When purified mitotic chromosomes were examined using fluorescence microscopy with DAPI staining, no contaminating interphase cells were detected (representative examples in Supplemental Fig. S1).

We performed 5C with a pool of primers targeting each end of each restriction fragment (a “double” alternating design (Hnisz et al. 2016)), which produces a complete interaction map for all restriction fragments throughout the two 2MB regions (Supplemental Fig. 2). In interphase cells, TADs are readily detected as domains of increased interaction frequencies between loci flanked by CTCF-bound sites (Fig. 1A, representative TAD marked with dashed line). By evaluating the insulation profile along the locus, we can identify TAD boundaries and quantify the strength of TADs (Gibcus et al. 2018; Crane et al. 2015). Insulation score is low at TAD boundaries and high at loci inside TADs (Fig. 1D, an example of a TAD is indicated with white dotted lines in Fig. 1A). As has been shown before, TAD boundaries are enriched in CTCF binding (The ENCODE Project Consortium 2012; Dixon et al. 2012; Rao et al. 2014; Vietri Rudan et al. 2015). Several CTCF-looping interactions are detected by their appearance as “dots” of elevated interaction frequency ((Fig. 1A), several representative loops marked with arrows). To illustrate CTCF-loops in another way, we plotted the interaction frequencies of one representative CTCF site with flanking loci (Fig. 1G). We find that interaction frequencies generally decay with genomic distance, but that peaks appear at other CTCF sites, consistent with loop formation. It has been shown in Hi-C, 4C and ChIA-PET datasets that loops between CTCF sites typically occur between motifs that are orientated convergent to each other (Rao et al. 2014; Wit et al. 2015; Tang et al. 2015; Guo et al. 2015). Looping interactions we observe in our 5C data are consistent with this (Fig. 1G).

**Figure 1.**
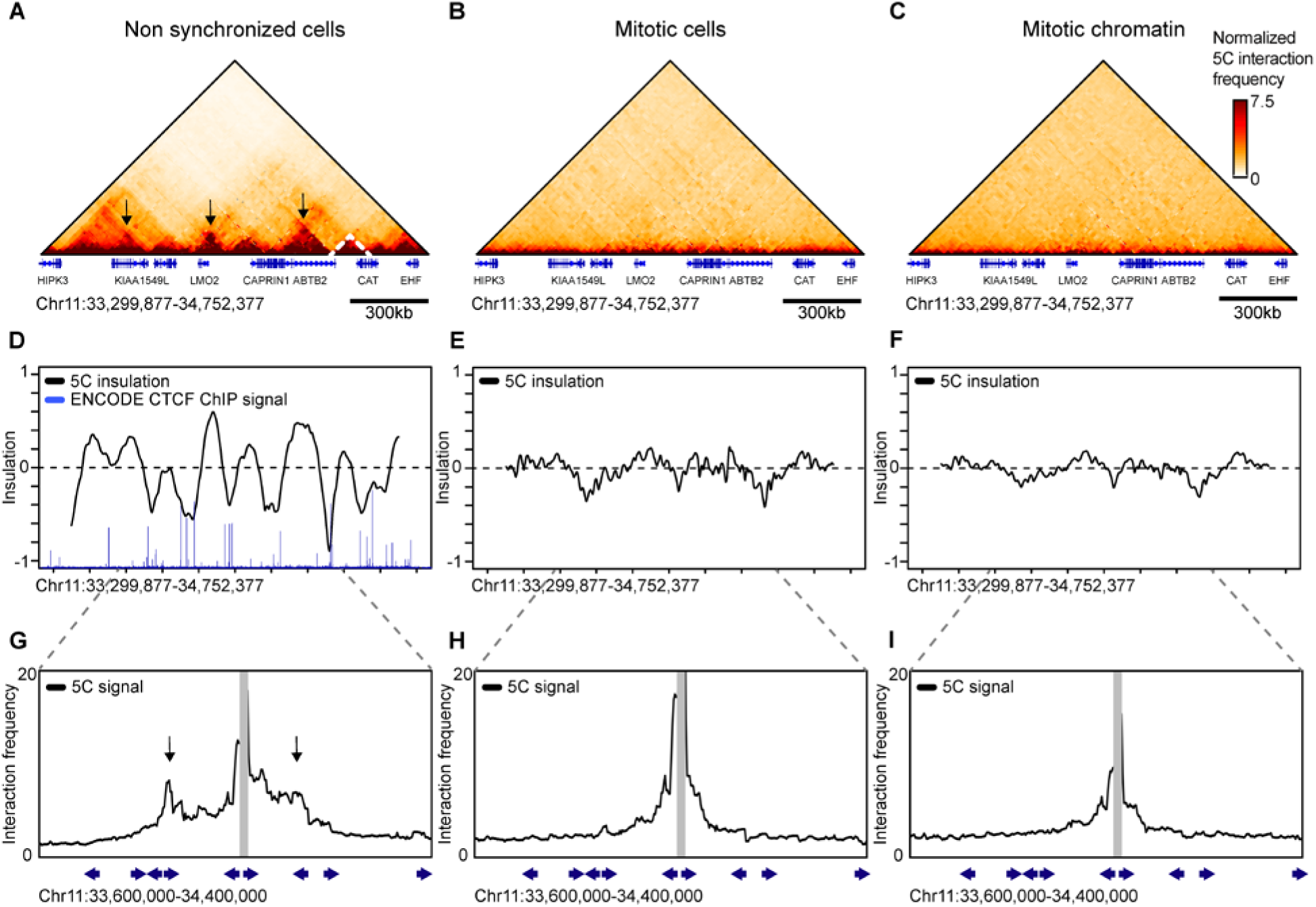
Topologically associating domains (TADs) and CTCF-loops are lost in prometaphase. 5C data of chromosome 11 33,299,877-34,752,377 shows TAD (marked by dashed lines) and CTCF loops (highlighted by arrows) in interphase **(A)**, however these structures are lost in nocodazole arrested mitotic cells **(B)** and purified mitotic chromatin **(C)**. 5C interaction data was used to calculate insulation profiles along the locus. Minima in insulations profiles represent TAD boundaries (**D-F**). The insulation profile for non-synchronized cells show a strong pattern alternating peaks centered within TADs and valleys where TAD boundaries are located (**D**). TAD boundaries are highly co-localized with bound CTCF as is shown by ENCODE CTCF ChIP-seq signal. Insolation profiles for mitotic cells and mitotic chromatin do not show deep minima indicating TAD boundaries are absent or strongly reduced (**E-F**). 5C interactions profiles anchored on one CTCF-bound site (15 kb bin spanning chr11: 34,012,377-34,027,377). Peaks along these profiles (arrows) indicate CTCF-loops observed in interphase (**G**). CTCF loops are not detected in mitotic cells and mitotic chromatin (**H-I**). Blue arrows underneath the x-axis represent the position and orientation of CTCF motifs.

5C analysis of mitotic cells and purified mitotic chromosomes reveals that TADs are no longer observed in mitotic cells and mitotic chromatin (Fig. 1B-C), in agreement with previous observations (Naumova et al. 2013; Gibcus et al. 2018). The loss of TADs can be seen both visually in the interaction heatmap, as well as by calculation of the insulation profile (Fig. 1E-F). Importantly, looping interactions between CTCF sites are also no longer detected in mitosis (Fig. 1H-I).

### Analysis of chromatin accessibility in interphase and prometaphase

To further investigate chromatin characteristics at CTCF binding sites, we determined chromatin accessibility using ATAC-seq (Buenrostro et al. 2015). ATAC-seq makes use of the Tn5 transposase to cut accessible chromatin and insert a sequencing adaptor. When two transposase events occur within 2 kb an ATAC-seq fragment is produced that can be amplified and sequenced. These fragments capture information in several different ways. First, ATAC-seq data provides information about genome-wide nucleosome positioning and spacing. This information is captured in the length distribution of all fragments (Fig. 2A). Very short fragments ranging from 24-80 bp are frequently observed. These fragments represent accessible regions in between nucleosomes or in between a nucleosome and another chromatin bound protein. Larger fragments typically form an enrichment of sizes that are multiples of approximately 195 bp. This reflects the nucleosomal array, which has been seen before in previous studies using ATAC-seq (Buenrostro et al. 2013). In mitotic cells and mitotic chromatin we observed a similar nucleosomal array. However, interestingly, the nucleosomal array in mitotic cells and chromatin is more pronounced, as the enrichment for fragments that are multiples of 195 bp, represented by the peaks in Fig. 2A, is stronger. This enrichment suggests that the spacing of nucleosomes is more regular in mitosis. It has been shown that histone H1 and other chromatin binding proteins are evicted during mitosis (Martinez-Balbas et al. 1995; Krishnan et al. 2017), which could result in a more regularly spaced array like we observe in the ATAC-seq data.

**Figure 2.**
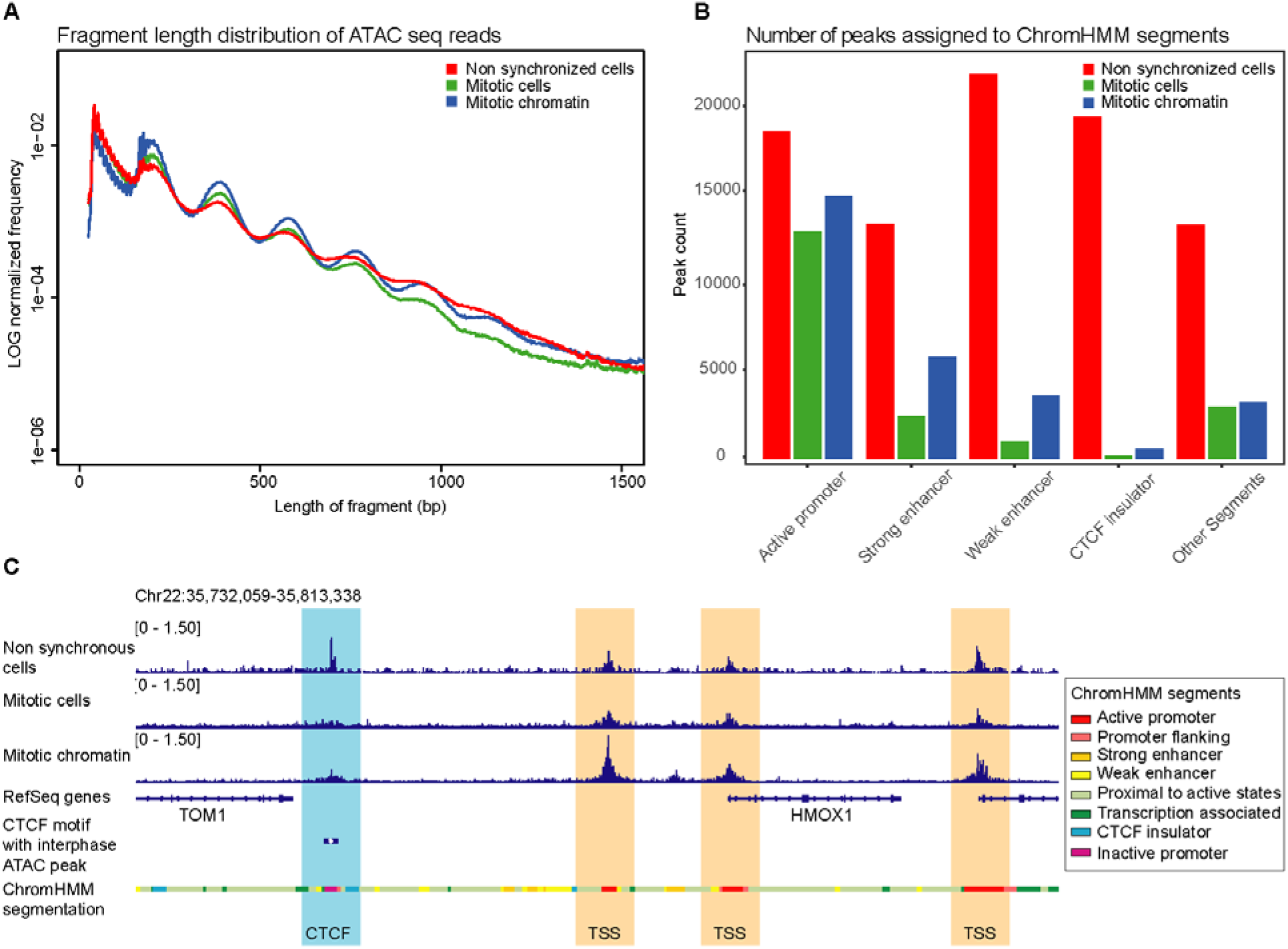
ATAC-seq data shows accessibility at active CTCF sites is reduced in mitosis, though maintained in other functional elements, such as TSSs. **(A)** Fragment length distribution of ATAC-seq reads genome-wide in non-synchronized cells, mitotic cells and purified mitotic chromatin. **(B)** Distribution of number of peaks called in non-synchronized cells, mitotic cells and mitotic chromatin and their position on ChromHMM segments. **(C)** Example of a representative region illustrating maintenance of accessible chromatin at TSSs in mitotic conditions while ATAC-seq signal is lost at CTCF motifs.

A second type of information captured in ATAC-seq data is on accessibility of specific classes of sites. We called peaks on ATAC-seq data using the HOMER software (Heinz et al. 2010). ATAC-seq data have been shown to display peaks in regions of open chromatin similar to peaks generated by DNase I sensitivity assays (Buenrostro et al. 2013). We can separate regions of the genome based on their function assigned by ChromHMM (Ernst and Kellis 2017; The ENCODE Project Consortium 2012). ATAC-seq peaks are found in different types of functional elements, such as active promoters, enhancers or CTCF insulator regions (Fig. 2B). We then compared peaks found in interphase to peaks found in mitotic cells and mitotic chromatin. In general, there is large loss of peaks in prometaphase compared to interphase. However, when we compare which types of functional elements lose peaks, we see that active promoters maintain significant accessibility, whereas the other functional elements examined lose most accessibility in mitosis (Fig. 2B). The maintenance of significant accessibility of promoter regions and concomitant loss of accessibility at enhancers during mitosis has been previously observed using DNase I sensitivity assays (Hsiung et al. 2015a; Martinez-Balbas et al. 1995). However, in addition to this, we also find an even more substantial loss of accessibility at CTCF binding sites in mitosis. This was observed in both mitotic (prometaphase) cells and mitotic chromatin isolated from prometaphase cells. Fig. 2C shows an example of ATAC-seq signal at individual TSSs and CTCF motifs (Fig. 2C).

### V-plots reveal protein occupancy at CTCF motifs and nucleosome positioning in interphase

In addition to determining the general accessibility of a region, ATAC-seq captures information on protein binding footprints and nucleosome positioning flanking hypersensitive sites. This information is captured by the length of fragments at a site of interest. We used V-plots to represent this data. V-plots have been used to plot MNase data as a way to display chromatin binding by site specific factors and positioning of flanking nucleosomes on different length scales (Zentner and Henikoff 2012). We have plotted our ATAC-seq data with the fragment length on the y-axis and positioning of the midpoint of the fragment on the x-axis representing the distance to the binding site of interest. V-plots can be produced for any regulatory element. In order to investigate local chromatin state at and around sites of CTCF binding, we made V-plots of our data on CTCF motifs (Kim et al. 2007) that are accessible, i.e. have an ATAC-seq peak, in interphase (Fig. 3A). In non-synchronized cells, we observe an enrichment of 80-100 bp fragment at the CTCF binding sites (see asterisk), which represents the footprint of bound CTCF protein and possibly associated proteins like cohesin. Similar footprints have been found using MNase digestion (Fu et al. 2008). The CTCF footprint can also be observed when the fragment length distribution is plotted for all fragments with their midpoint on bound CTCF motifs (Fig. 3D, marked by asterisk). When we plot the lengths of reads with one end on a CTCF motif, we observe that many fragments have a short length (Fig. 3E). These represent fragments generated by pairs of ATAC cleavages in between bound CTCF and the flanking nucleosomes.

**Figure 3.**
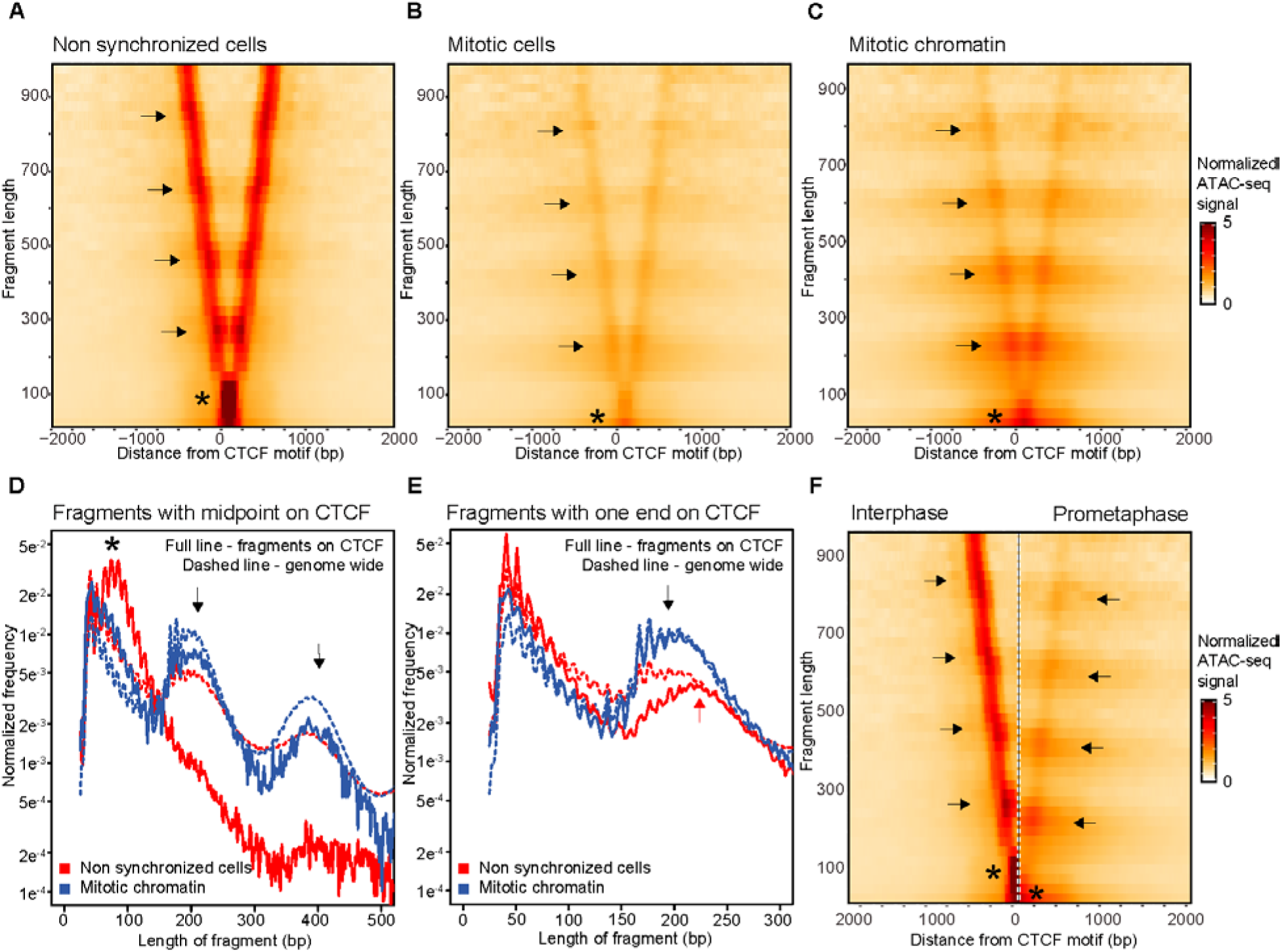
ATAC-seq data represented in V-plots show a loss of CTCF binding in mitosis and a rearrangement of nucleosomes flanking CTCF motifs. (A-C) ATAC-seq data represented in V-plots aggregated at CTCF sites. The lengths of ATAC-seg reads are plotted on the y-axis, and the distance between their midpoints and the CTCF motif is shown on the x-axis. **(A)** V-plot for interphase ATAC-seq data shows enrichment of 80-100 bp fragments centered at the CTCF motif (asterisk), representing a CTCF footprint. Enriched dots along the arms of the V (arrows) represent strongly positioned nucleosomes. (B) and (C): V-plots for ATAC-seq data mitotic cells isolated mitotic chromatin **(C)** aggregated at CTCF motifs with interphase ATAC-seq peaks. The enrichment for 80-100 bp fragments centered at CTCF motis is lost, enriched dots along the arms of the V are positioned at smaller lengths and a weak horizontal banding pattern is observed. **(D)** Distribution of fragment lengths of reads that have their midpoint on a CTCF motif. Dased line represents the genome-wide average read length distribution. Arrows indicate read lengths representing 1 and 2 nucleosomes. **(E)** Distribution of fragment length of reads with either read end near a CTCF motif with interphase ATAC-seq peak compared to the genome-wide average (dashed line). In interphase reads representing one flanking nucleosome are longer (red arrow) as compared to reads representing one flanking nucleosome in mitosis (black arrow). **(F)** Side-by-side comparison of V-plots for both non-synchronized and mitotic chromatin illustrate the shift in size at which ATAC-seq fragments are enriched along the arms of the V. The shift in nucleosome positioning is highlighted using arrows. Highlighted using asterisks are loss of the CTCF footprint in mitosis compared to interphase.

The second type of information that can be derived from V-plots is regarding the positioning of flanking nucleosomes. For bound CTCF motifs in interphase, we observe enriched dots on the arms of the V in the V-plot (Fig. 3A, marked by arrows). These dots indicate strong positioning of several nucleosomes flanking the bound CTCF motif, consistent with previous MNase results (Fu et al. 2008). The series of enriched signals represent ATAC-seq fragments covering one, two, three and four flanking nucleosomes, but are longer than expected for a typical nucleosomal array. We attribute this size discrepancy to mean that some of these fragments can cover not only one or more nucleosomes, but also the flanking bound CTCF site. This becomes even clearer when fragment lengths are plotted of reads which have one of their read ends near a bound CTCF motif (Fig. 3E, marked with red arrow). We observe an enrichment of fragments that are around 220 bp, instead of the expected 195 bp for a canonical mononucleosome. Similar results were found when V-plots were made for all CTCF motifs or for CTCF motifs with peaks from available CTCF ChIP-seq ENCODE data produced by the Bernstein lab (The ENCODE Project Consortium 2012) (Supplemental Fig. S3A). Taken together, these analyses show that during interphase CTCF is bound to its motif and nucleosomes flanking CTCF-occupied sites form a regularly spaced array.

### V-plots show loss of binding at CTCF motifs and rearrangement of nucleosomes in prometaphase

We then created V-plots for ATAC-seq data generated from mitotic cells and purified mitotic chromatin to analyze the CTCF footprint and the positioning of the flanking nucleosomes at CTCF motifs that have an interphase ATAC-seq peak (Fig. 3B-C). As expected from the peak calling assessment, overall accessibility is reduced (and no longer significant) in mitotic conditions. However, some accessibility remains. We do note that ATAC-seq signal at CTCF sites for purified mitotic chromatin is stronger compared to ATAC-seq signal in mitotic cells. This higher signal to noise ratio is possibly caused by lower background signals overall due to stronger nucleosome interactions as a result of detergents in the chromosome purification buffer (Gasser and Laemmli 1987). Several additional features are observed in V-plots of CTCF motifs in mitotic conditions. First, there is a change in the CTCF footprint in mitosis. The enrichment of 80-100 bp fragment lengths observed in interphase is no longer present. Instead there is an enrichment of small fragments around 25-75 bp (marked by asterisks). This suggests that CTCF is no longer bound to the motif, where it protected the site from ATAC-seq cleavage in interphase. We observed this in both mitotic (prometaphase) cells and mitotic chromatin purified from prometaphase arrested cells. The loss of 80-100 bp fragments is also observed when the length distribution is plotted for fragments with their midpoint on interphase bound CTCF motifs (Fig. 3D, compare to interphase length distribution marked by asterisk).

Second, the positioning of nucleosomes around interphase bound CTCF motifs also changes during mitosis (Fig. 3B-C, marked by arrows). Loss of CTCF binding in mitosis would create a large accessible region between the flanking nucleosomes, which can in turn cause nucleosomes to move inwards. This movement of nucleosomes would cause the linkers in the nucleosomal array to become larger than average. We observe this phenomenon in several ways in our V-plots. First, nucleosomes are able to occupy CTCF sites in mitosis. This can be seen by a gain of mono- and dinucleosome sized fragments in the length distribution plot in Fig. 3D (marked by arrows). Nucleosome occupancy at the CTCF motif is not observed in interphase when CTCF typically occupies the motif. Second, as discussed above, in interphase we observe a peak at 220 bp in the fragment length distribution for fragments with one end in a CTCF motif that is larger than typical for a mononucleosome (Fig. 3E, marked by red arrow). In mitosis, this mononucleosome peak becomes more similar to the genome-wide average (195 bp); again suggesting that CTCF is no longer bound (Fig. 3E, marked by black arrow). This size discrepancy becomes even more obvious when V-plots for non-synchronized cells and purified mitotic chromatin are plotted side by side (Fig. 3F, compare arrows). Lastly, there is a change in nucleosome spacing and positioning during mitosis (Fig. 3B-C). In interphase, the locations of nucleosomes are observed as enriched dots along the arms of the V with a strong positioning relative to the CTCF motif. Conversely, in mitosis, the enriched dots become less pronounced and we observe horizontal bands of elevated fragment frequency in the heatmap running several kb up and down stream of the CTCF motif. Given that these V-plots are normalized for genome-wide average fragment length frequency, a banding pattern only emerges when the spacing between several of the flanking nucleosomes differs and/or is more variable from the genome-wide average. Specifically, the banding pattern indicates that there is an increase in nucleosomes spaced by larger linkers than the genome-wide average (Fig. 3E, compare to dashed line). These observations confirm that several flanking nucleosomes are able to move inwards, creating longer linkers between them. This creates a local nucleosomal array around CTCF sites with larger than average linkers, which is also confirmed by general accessibility detected by short ATAC-seq products in a region from −500 to +500 bp from the CTCF motif.

Taken together, our data suggest that CTCF is no longer bound in prometaphase and nucleosomes rearrange. To exclude that there are certain subgroups of CTCF motifs that maintain binding in mitosis, we plotted V-plots for several obvious classes of CTCF motifs, e.g. motifs proximal or distal to TSSs or CTCF motifs that maintain an ATAC-seq peak in mitosis for fragment lengths of 75-150 bp at CTCF sites, using k-means clustering to determine whether there are subgroups of motifs that maintain a CTCF footprint in mitosis (Supplemental Fig. S4). None of these methods found a specific group of CTCF motifs that maintain CTCF binding, suggesting that binding to CTCF motifs is generally lost by late prometaphase. Finally, we confirmed the loss of CTCF binding and rearrangement of nucleosomes in multiple differentiated cell lines (Supplemental Fig. S5-6).

### Transcription start sites maintain significant accessibility, but show variable loss of factor binding and nucleosome repositioning in prometaphase

In addition to our findings regarding CTCF-bound motifs, we also made several important observations about the chromatin landscape at accessible TSSs in interphase and mitosis (Supplemental Fig. S7). First, we see an enrichment of fragments with a length below 150 bp compared to the genome-wide average in interphase (Supplemental Fig. S7A marked by asterisk). We do not observe an enrichment of fragments of a defined size as we do for CTCF (compare supplemental Fig. S7D to Fig. 3D). This can be explained by the fact that contrary to CTCF sites, TSSs and the local chromatin around TSSs are bound by multiple different proteins and which proteins are binding at individual TSSs is more diverse. Second, we do not observe enrichments of nucleosome sized fragments centered on the TSS (Supplemental Fig. S7D), confirming that in interphase typically no nucleosomes are bound at TSSs. Furthermore, the length of fragments covering the two nucleosomes directly flanking the TSS appears less defined compared to those flanking CTCF sites (compare Fig. 3E to Supplemental Fig. S7E, indicated with red arrow). There is a local enrichment of fragments a little over 200 bp in length. These fragments represent ATAC-seq products covering factors bound to the TSS plus the neighboring nucleosome, resulting in a length that is over the genome-wide expected 195 bp covering a single nucleosome. In addition, the enrichment of fragments over 200 bp is less pronounced compared to genome-wide average for single nucleosome fragments, suggesting that heterogeneity in factors binding to the TSS can result in a wider distribution of fragment lengths covering the TSS and the flanking nucleosome.

In mitotic conditions, complementary to our peak calling analysis (Fig. 2B-C), V-plots show that overall accessibility at TSSs is reduced in prometaphase, though maintained at higher levels than at CTCF sites (compare Fig. 3B-C to Supplemental Fig. S7B-C). We note that accessibility at TSSs, though reduced, is still significant (see Fig. 2B-C). Interestingly, we observe that in mitotic chromatin, nucleosome sized fragments are observed at the TSS (Supplemental Fig. S7D, marked by arrows), suggesting that nucleosomes start occupying TSSs in prometaphase. This led us to explore further how mitotic accessibility at TSSs is related to nucleosome positioning. We compared the accessibility and nucleosome positioning in the top 20 percent most accessible and 20 percent least accessible TSSs in mitosis (see methods). In interphase, these two groups behave highly similarly, except for signal intensity as expected (compare Fig. 4A and D to Fig. 4F and I). In mitosis, both groups of TSSs can be occupied by nucleosomes, however the level of nucleosome occupancy is inversely related to the level of remaining accessibility in mitosis (Fig. 4D and I, marked by black arrows). For the group of TSSs with low levels of remaining accessibility, the nucleosome occupancy in mitosis is comparable to the genome-wide average (Fig. 4D, compare to dotted line), while the group of TSSs with the highest remaining accessibility in mitosis, the nucleosome occupancy is lower than the genome-wide average (Fig. 4I, compare to dotted line). During mitosis, the set of TSSs with the lowest accessibility resembles CTCF sites in several ways. First, while in interphase the fragments representing mononucleosomes flanking the TSS are longer than genome-wide average, in mitosis this length shortens to the canonical 195 bp (Fig. 4E, compare black and red arrows). This shortening of fragments covering flanking nucleosomes can also be observed in V-plots (Fig. 4C, compare arrows). Lastly, as for CTCF sites, we observe a gain of horizontal banding pattern in the V-plot, again indicating that there is an increase in nucleosomes spaced by larger linkers than the genome-wide average. Importantly and in contrast to CTCF sites, these TSSs are significantly more accessible than the genome-wide average, despite the fact that this set of TSSs is now occupied by nucleosomes (see also Fig. 2B-C).

**Figure 4.**
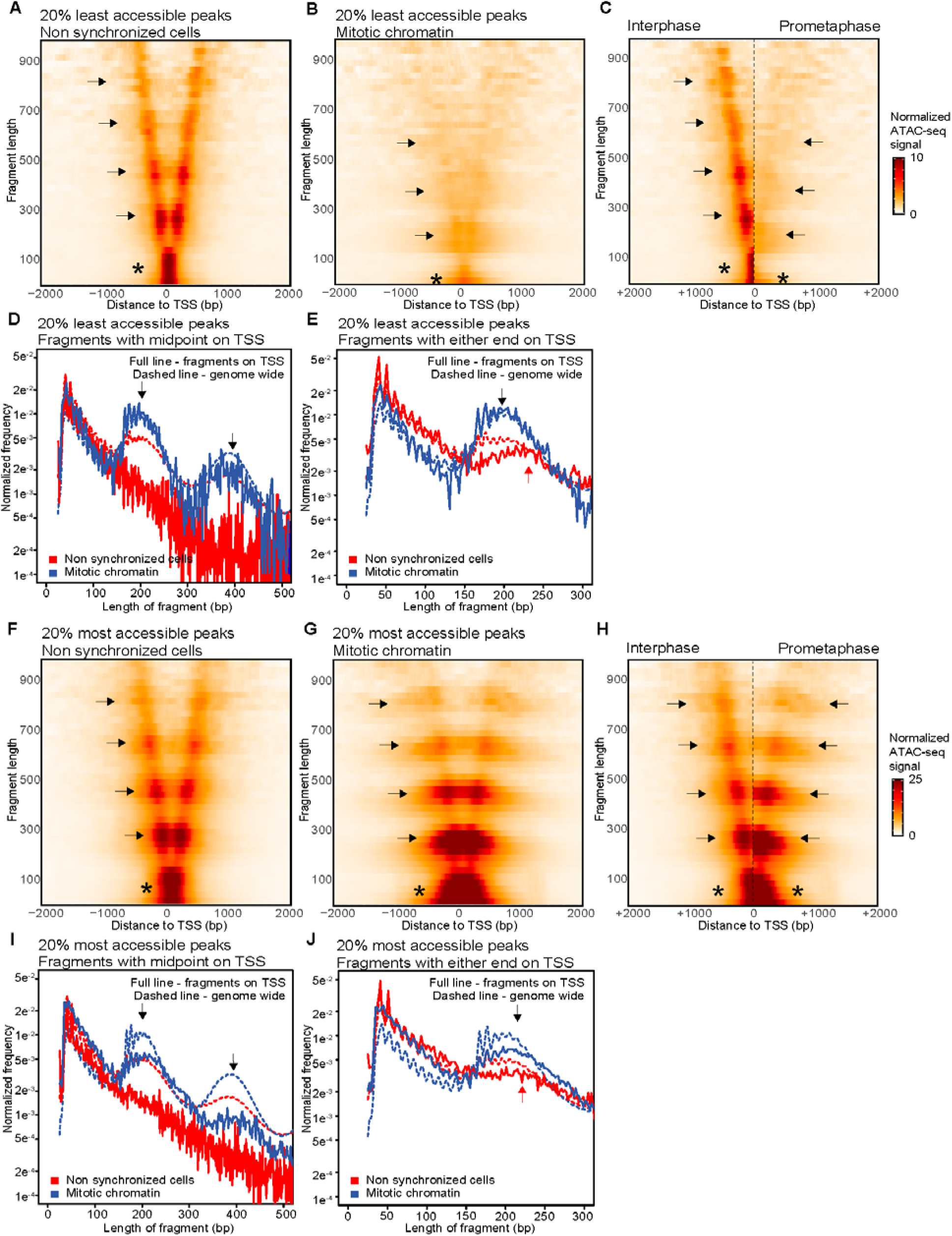
ATAC-seq data represented in V-plots shows nucleosome repositioning, while accessibility is maintained at transcription start sites. Panels A-E show ATAC-seq data for the set of TSSs that display the lowest 20 percent accessible peaks in mitotic chromatin, panels F-J show ATAC-seq data for the set of TSSs that display the highest 20 percent accessible peaks in mitotic chromatin **(A-B)** V-plots of ATAC-seq signal aggregated at TSSs in non-synchronized cells **(A)** and mitotic chromatin **(B)**. **(C)** Side-by-side comparison of the V-plots of 2kb upstream of low accessible TSSs for both non-synchronized cells and mitotic chromatin reveals a reduced accessibility overall, with reduced enrichment of dots along the arms of the V and a downwards shift of horizontal bands. **(D)** Distribution of fragment length of reads with their midpoint on a TSS compared to the genome-wide average (dashed line). Arrows indicate read lengths representing 1 and 2 nucleosomes. **(E)** Distribution of fragment length of reads with either end in a TSS in mitotic chromatin compared to the genome-wide average (dashed line). In interphase reads representing one flanking nucleosome are longer (red arrow) as compared to reads representing one flanking nucleosome in mitosis (black arrow). **(F-G)** V-plots of ATAC-seq signal aggregated at TSSs that display the highest 20% accessibility in mitotic chromatin for non-synchronized cells **(F)** and mitotic chromatin **(G)**. **(H)** Side-by-side comparison of the V-plots shown in **(F)** and **(G)** reveals a comparable accessibility overall, with enrichment of dots along the arms of the V at the same fragment length for both non-synchronized and mitotic chromatin. A horizontal banding pattern is stronger in mitotic chromatin. **(I)** Distribution of fragment length of reads with their midpoint on a TSS with highest 20% highest accessible peaks in mitotic chromatin compared to the genome-wide average (dashed line). Arrows indicate read lengths representing 1 and 2 nucleosomes. **(E)** Distribution of fragment length of reads with either end in a TSS with highest 20% highest accessible peaks in mitotic chromatin compared to the genome-wide average (dashed line). For mitotic chromatin reads representing one flanking nucleosome are of similar length (black arrow) as compared to reads representing one flanking nucleosome in interphase (red arrow).

The set of TSSs that maintain high levels of accessibility in mitosis however behaves differently. For this set of TSSs we continue to observe an enrichment of short fragments (below 150 bp) compared to the genome-wide average and larger fragments covering the flanking nucleosomes (Fig. 4J, marked by arrows), suggesting that some factors remain bound to the TSSs. This is also observed by a lack of downwards shift of nucleosome sized fragments along the arms in the V (Fig. 4H, compare arrows). However, as indicated above, some increase in nucleosome occupancy at the TSS is observed in mitosis, suggesting that this set of TSSs is heterogeneous with some sites gaining nucleosomes, while other sites maintain interphase nucleosome positioning. The presence of a subset of TSSs with repositioned nucleosomes in mitosis is also reflected in the gain of horizontal bands in the V-plot (Fig. 4H). As discussed above, the horizontal banding pattern is a result of repositioned nucleosomes that have larger and more variable linker lengths than the genome-wide average.

### CUT&RUN directly observes loss of CTCF binding in prometaphase

Next, we set out to detect CTCF binding directly. Given concerns about formaldehyde induced artifacts that affect protein binding to mitotic chromosomes (discussed above, and (Pallier et al. 2003; Teves et al. 2016)), we decided not to use ChIP-seq. Instead, we use CUT&RUN (Skene and Henikoff 2017). In contrast to ChIP-seq, CUT&RUN can be done on unfixed cells and targets MNase to sites where the protein of interest is binding using primary antibodies and a protein A-MNase fusion. An additional benefit of CUT&RUN is that it does not require any pull down of molecules. CUT&RUN signal for CTCF shows a very clear peak at accessible CTCF motifs in interphase. However, in mitosis, the signal diminishes to almost background levels (Fig. 5A). Increasing MNase digestion time makes it possible to detect weaker bound proteins. We tested several digestion times for both interphase and mitotic cells (Supplemental Fig. S8). Even prolonged digestion (30 minutes) did not reveal CTCF binding in mitotic (prometaphase) cells. We analyzed CTCF binding in different obvious classes of CTCF motifs, e.g. sites with an ENCODE CTCF ChIP-seq peak or sites proximal to TSSs (Supplemental Fig. S9). None of these sites showed CTCF CUT&RUN signal in prometaphase cells. Furthermore, when we represent CTCF CUT&RUN signal of each motif sorted on signal strength, there are no CTCF sites found that maintain CTCF binding (Supplemental Fig. S10). This suggests that CTCF binding does not just become weaker in prometaphase, but that essentially all motif-specific binding is lost.

**Figure 5.**
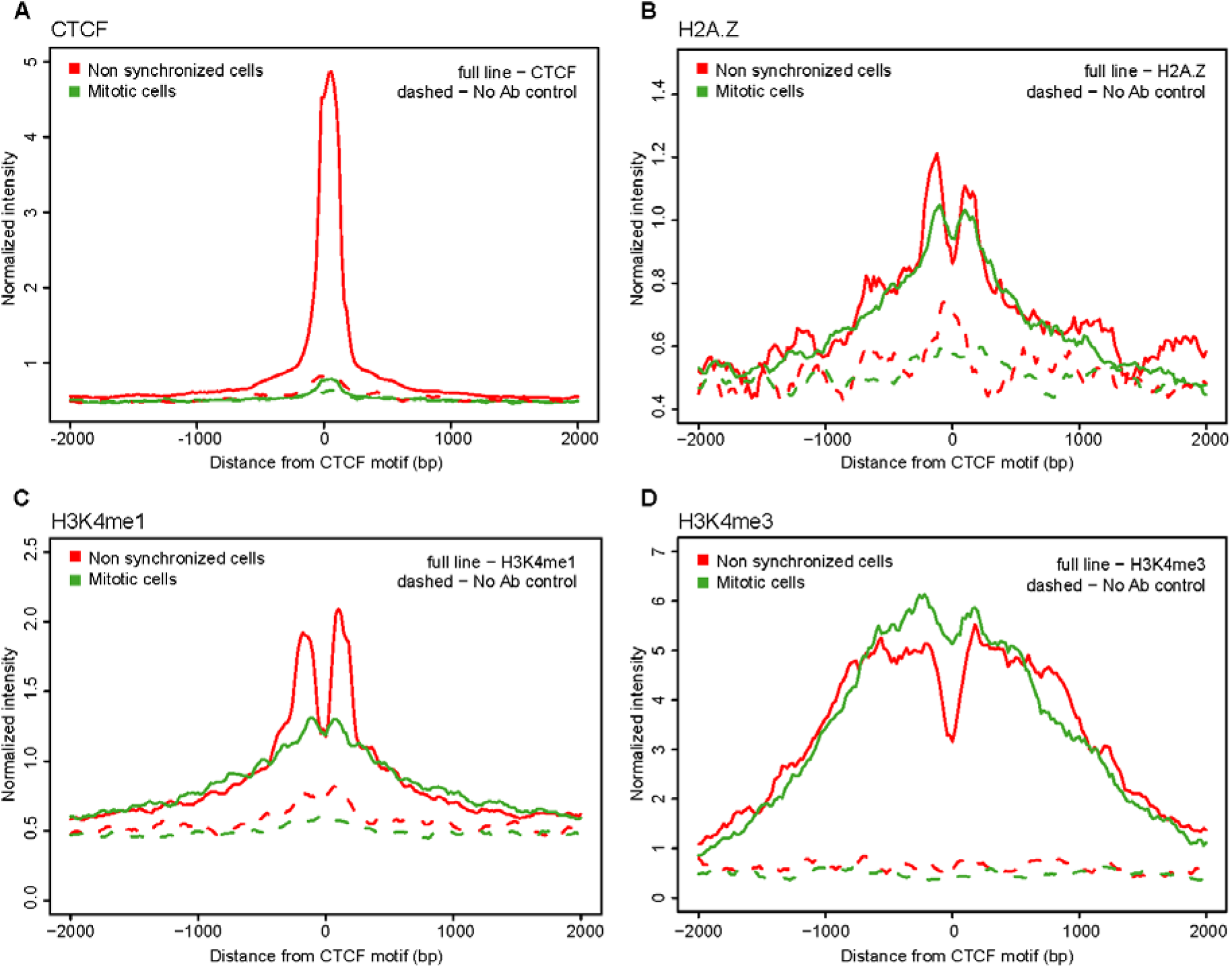
CUT&RUN data shows loss of CTCF binding at CTCF motifs in mitosis, while H2A.Z and H3K4 methylation marks are maintained. **(A)** Aggregation plot of CTCF CUT&RUN signal of reads shorter than 120 bp after 10 min digestion. Reads are aggregated on CTCF motifs that display an interphase ATAC-seq peak. **(B)** Aggregation plot of H2A.Z CUT&RUN signal of reads longer than 120 bp after 30 min digestion. Reads are aggregated at CTCF motifs with interphase ATAC-seq peak. **(C)** Aggregation plot of H3K4me3 CUT&RUN signal of reads longer than 120 bp after 30 min digestion on CTCF motifs with interphase ATAC-seq peak. **(D)** Aggregation plot of H3K4me3 CUT&RUN signal of reads longer than 120 bp after 30 min digestion on CTCF motifs with interphase ATAC-seq peak.

Next, we wanted to determine whether CTCF gains binding at other sites, while it is losing binding at the described CTCF motif. Whereas de novo peak calling of CTCF CUT&RUN data in interphase identified 7824 peaks, in mitotic cells only 107 peaks were identified (Supplemental Fig. S11). This suggests there is very little site specific binding in mitosis. It has been described that CTCF becomes highly phosphorylated in mitosis (Dephoure et al. 2008). To ensure the loss of CUT&RUN signal is not caused by the inability of the CTCF antibody to recognize phosphorylated CTCF, we performed western blot with lysates of non-synchronized cells, mitotic cells and mitotic chromatin (Supplemental Fig. S12). The antibody detected CTCF in both non-synchronized and mitotic cell extracts. Interestingly, we did not detect CTCF in extracts from mitotic chromatin (Supplemental Fig. S12). This indicates again that CTCF binding in mitosis is lost and when isolating chromatin from prometaphase arrested cells, CTCF is not co-purified with the chromatin.

### Histone marks and variants at active CTCF sites are maintained in mitosis

There are several histone characteristics described to date that are associated with CTCF bound motifs in interphase. The histone variant H2A.Z is often observed on nucleosomes flanking CTCF sites as well as to TSSs (Fu et al. 2008; Kelly et al. 2010; Nekrasov et al. 2012). Additionally, several histone modifications have been found to co-localize with bound CTCF motifs; for example H3K4me1 for sites distal to TSSs and H3K4me3 for sites proximal to TSSs (Fu et al. 2008). In order to assay the presence of these histone marks and variants, we performed CUT&RUN. For H2A.Z and H3K4 mono- and trimethylation we observe two distinct peaks around the CTCF motif in non-synchronized cells (Fig. 5B-D). This confirms our observations in V-plots of ATAC-seq data, where the CTCF flanking nucleosomes are strongly positioned relative to distance from the CTCF motif in interphase due to CTCF binding.

Many histone marks and variants have been suggested to serve as bookmarks for factor binding in mitosis (reviewed in (Wang and Higgins 2013)). Both H2A.Z and H3K4 methylation states have been observed throughout the cell cycle (Nekrasov et al. 2012; Kelly et al. 2010; Liu et al. 2017; Javasky et al. 2017). Using CUT&RUN we observed that H2A.Z binding and H3K4 methylation states are maintained at CTCF sites during mitosis, even though CTCF binding is temporarily lost (Fig. 5B-D). We note that the level of H3K4me1 is reduced during mitosis, in contrast to H3K4me3 for which signal strength is similar in interphase and mitosis. It has been previously shown that H3K4me3 typically gets rapidly reestablished on histones on both sister chromatids in S-phase, while H3K4me1 levels do not get restored until after cell division (Lin et al. 2016). A 50% reduction in H3K4me1 levels as a result of dilution over the two sister chromatids is consistent with the reduced levels observed by CUT&RUN. Furthermore, as described, we observe two strong peaks in H2A.Z and H3K4 methylation signal flanking the CTCF motif in interphase, however in mitosis these peaks become weaker and form two broader peaks with a less defined valley in between. This is in concordance with ATAC-seq data, where we observed rearrangement of the nucleosomes and filling in of the nucleosome depleted regions (NDRs) at CTCF in mitosis.

We then analyzed H2A.Z and H3K4me3 levels at TSSs detected by CUT&RUN in interphase and mitosis. In line with previous studies and our CUT&RUN data at and around CTCF sites, we found that H2A.Z and H3K4me3 levels are maintained at TSSs in prometaphase (Supplemental Fig. S13; (Lin et al. 2016; Varier et al. 2010; Wang and Higgins 2013; Nekrasov et al. 2012; Kelly et al. 2010; Javasky et al. 2017)). Interestingly, but not surprisingly, we again observe loss of depletion of histone signal at TSSs in prometaphase, similar to CTCF sites. This is an indication that nucleosomes reposition and occupy at least a subset of TSSs, consistent with our observations in ATAC-seq V-plots (Fig. 5).

### Live cell imaging shows that CTCF is not enriched on prometaphase chromosomes and that essentially all specific binding is lost

Genomic and imaging studies have occasionally reported contradicting findings when it comes to factor binding to prometaphase chromatin. To independently validate our observation using genomics techniques that essentially all site specific CTCF binding is lost during prometaphase, we turned to live-cell imaging and applied a three-pronged approach based on long-term time-lapse imaging, Fluorescence Recovery After Photobleaching (FRAP) and Single-Particle Tracking (SPT). We previously generated and validated a U2OS cell line where all CTCF alleles have been Halo-tagged (C32 Halo-CTCF; (Hansen et al. 2017)). To visualize mitotic cells, we additionally stably integrated a histone H2B-GFP transgene. We performed ATAC-seq on this cell line and observed the expected loss of accessibility at CTCF sites in mitosis and loss of the CTCF footprint represented in V-plots (Supplemental Fig. S5E-F and Supplemental Fig. S6G-H). First, we used multi-hour time-lapse fluorescence microscopy to observe Halo-CTCF (Movie S1-2) and H2B-GFP (Movie S2) in actively dividing cells. Although CTCF was clearly enriched on mitotic chromosomes during most phases of mitosis (e.g. telophase), CTCF localization appeared to be diffuse during prometaphase. Second, to observe CTCF at higher resolution and quantify its binding dynamics, we used FRAP. As for the genomics experiments, we used nocodazole to arrest cells in prometaphase. As we observed with time-lapse microscopy, CTCF showed a diffuse localization without clear enrichment on mitotic chromosomes during prometaphase (Fig. 6A; upper panel). To rule out any artifacts due to nocodazole drug treatment, we also identified cells in prometaphase without drug-treatment based on their H2B-GFP localization (“prometaphase enriched”) and similarly observed diffuse CTCF localization without enrichment on chromatin.

**Figure 6.**
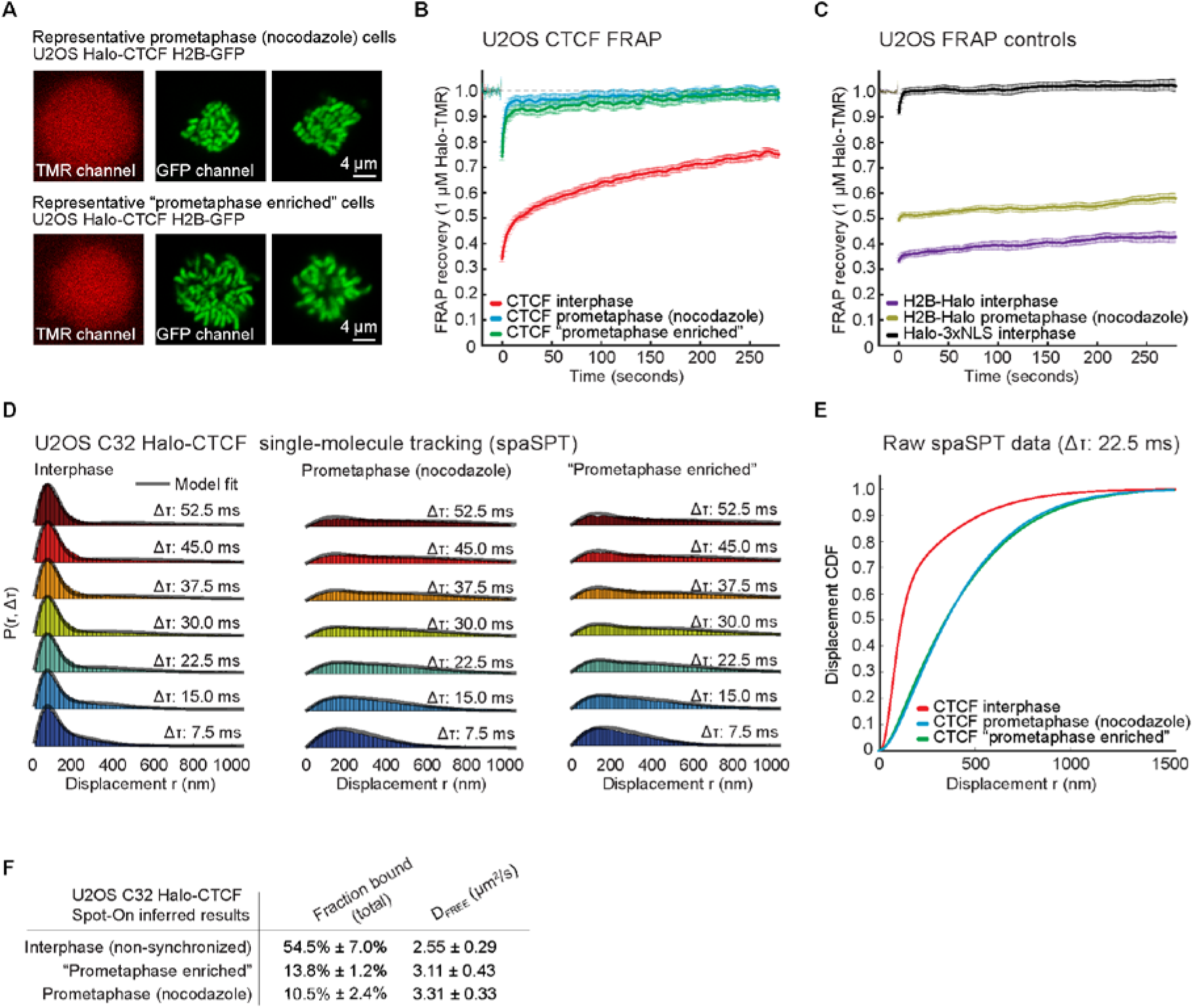
Live cell imaging techniques show that CTCF is more dynamically bound and that the fraction of CTCF bound to chromatin is reduced in mitosis. **(A)** H2B-GFP localization of representative U2OS cells for prometaphase arrested cells and non-arrested cells selected as prometaphase (“prometaphase enriched”). **(B)** Fluorescence recovery after photo bleaching (FRAP) for Halo-CTCF in interphase, prometaphase arrested and prometaphase enriched cells. **(C)** Controls showing FRAP for H2B-Halo in interphase and prometaphase arrested cells and for Halo-3xNLS in interphase cells. **(D)** Single particle tracking displacement statistics for halo-CTCF in different timeframes in interphase, prometaphase arrested and prometaphase enriched cells. **(E)** Displacement cumulative distribution function (CDFs) derived from single particle tracking (SPT) at Δτ 22.5ms for interphase, prometaphase arrested and prometaphase enriched cells. **(F)** Fraction bound of Halo-CTCF calculated using the Spot-On model in interphase, prometaphase arrested and prometaphase enriched cells (Hansen et al. 2018).

We then performed FRAP, bleaching a ~1 μm circle and quantifying the recovery (Hansen et al. 2017). In interphase, Halo-CTCF showed slow recovery consistent with a high fraction of chromatin binding to specific CTCF sites with an apparent residence time of a few minutes (Fig. 6B; (Hansen et al. 2017)). However, in prometaphase, Halo-CTCF showed ~90-95% recovery within seconds as well as a fraction of the population (~5-10%) which showed slower recovery suggesting stable binding by only a small subpopulation (Fig. 6B). We validated our FRAP approach and show that the difference in recovery was not due to improper drift-correction using histone H2B controls (Fig. 6C; note that the H2B bleach depth is slightly lower in prometaphase due to “gaps” between chromosomes, but that the rate of recovery is unchanged). Thus, in prometaphase, although a small population approaching our detection limit (~5-10%) does appear to bind specific sites with a residence time in the minute range, the vast majority of specific CTCF binding is clearly lost, which is consistent with the genomics experiments. We conclude that nearly all specific CTCF binding is lost in prometaphase.

Third and finally, we sought to verify this using an independent technique. We therefore turned to stroboscopic photo-activation single-particle tracking (spaSPT). spaSPT makes it possible to observe single CTCF molecules in live cells and to visualize both bound (specific and non-specific) and freely diffusing molecules without motion-blur bias (Hansen et al. 2018, 2017). We then tracked single CTCF molecules at 134 Hz in interphase (Movie S3), in nocodazole-arrested prometaphase (Movie S4) and in “prometaphase-enriched” cells (Movie S5) and quantified the distribution of displacements between frames. Analysis of the cumulative distribution function (CDF) revealed that nearly all CTCF chromatin binding was lost in prometaphase (Fig. 6D-E; bound molecules typically appear below 150nm). To quantitatively analyze the spaSPT data we fit a 2-state kinetic model (Spot-On) wherein CTCF can exist in either a bound (both specific and transient non-specific binding) and free state (Fig. 6D; (Hansen et al. 2018, 2017)). This revealed that whereas ~55% of CTCF molecules are chromatin bound in interphase, only ~10-14% are bound in prometaphase (Fig. 6F). Since this bound population includes transient non-specific binding, these results independently confirm the FRAP results (Fig. 6B). In summary, our live-cell imaging results independently confirm that nearly all specific CTCF binding is lost during prometaphase and, additionally, that this observation is not an artifact of nocodazole treatment.

## Discussion

Our study presents a comprehensive analysis of CTCF binding and CTCF motif-flanking nucleosomes in mitosis using several different complimentary techniques and using multiple differentiated cell lines. Genomics and live cell imaging techniques used here determine that overall CTCF binding is lost in mitosis, as are TADs and loops between CTCF sites. We did not find any subgroup of CTCF sites that maintain binding in mitosis, nor did we find any new sites of mitotic CTCF binding. This could explain why TADs and CTCF loops are not observed in mitosis. In addition to this, we found that nucleosomes flanking interphase CTCF sites rearrange in prometaphase, resulting in nucleosomes occupying the CTCF motif and forming an array of nucleosomes with larger and more variable linkers (Fig. 7). Furthermore, similar phenomena can occur at TSSs, although these sites remain hyperaccessible as observed by ATAC-seq. Epigenetic marks, such as histone variant H2A.Z and H3K4 methylation marks, are maintained at both CTCF sites and TSSs in mitosis.

**Figure 7.**
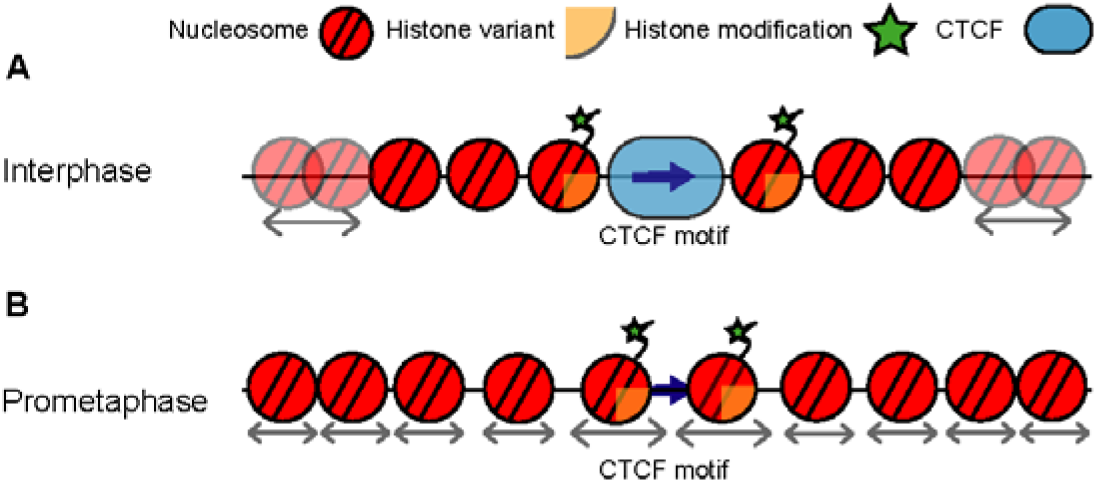
Dynamics of CTCF binding and chromatin organization around active CTCF motifs throughout the cell cycle. (**A**) In interphase CTCF is bound to its motif and flanking nucleosomes are strongly positioned relatively to the motif. Flanking nucleosomes are characterized by histone variants and modifications. (**B**) In prometaphase, CTCF binding is temporarily lost and nucleosomes rearrange to fill in the nucleosome depleted region at the CTCF motif. This increases the linker lengths between adjacent nucleosomes. Epigenetic marks however are maintained, possible functioning as bookmarks that enable inheritance of active CTCF motifs throughout the cell cycle. Arrows underneath nucleosomes indicate that the position of these nucleosomes can vary between cells.

Previous studies found evidence for CTCF binding to mitotic chromosomes using imaging and chromatin fractionation approaches (Cai et al. 2018; Liu et al. 2017; Burke et al. 2005). Additionally, proteomics studies of isolated mitotic chromatin detect CTCF, although at reduced levels compared to interphase chromatin (Gibcus et al. 2018; Ohta et al. 2010). Importantly however, all these approaches measure general mitotic chromatin association and do not capture information on site specific binding (Raccaud and Suter 2018). Our live cell imaging data also indicate that CTCF remains associated with chromatin during several stages of mitosis, however in prometaphase CTCF binding dynamics are changed and the vast majority of specific and stable binding is lost. This is complementary to our findings using genomics techniques, where we also observe loss of CTCF binding at interphase sites and we do not find any mitotic site specific binding. It is possible that CTCF remains associated with mitotic chromatin, although in a non-specific and highly dynamic manner. First, mitotic chromatin retention could enable proper segregation of CTCF levels over the daughter cells. Second, maintained chromatin association can enable efficient reestablishment of CTCF binding upon mitotic exit. A recent study observed a rapid raise of CTCF levels associated to the chromatin in late anaphase, as for many other chromatin binding factors (Cai et al. 2018). Additionally, we note that CTCF may possibly show cell type specific dynamics in prometaphase. Our study observes CTCF cell cycle dynamics of differentiated cells, using both transformed and non-transformed cell lines.

Transcription start sites are highly accessible and free of nucleosomes in non-synchronized cells that are mostly in interphase. In contrast to CTCF sites, TSSs remain hyperaccessible during mitosis. This has also been observed by DNAse I sensitivity assays (Hsiung et al. 2015b; Martinez-Balbas et al. 1995). Here we find that despite remaining highly accessible, nucleosome-sized ATAC-seq fragments are detected at TSSs in mitosis indicating that nucleosomes are able to occupy these sites, and that the spacing between flanking nucleosomes becomes larger and more variable compared to genome-wide average, similar to what we observed at and around CTCF sites. These observations are consistent with a recent study that found that a large fraction of NDRs at TSSs become filled in by a nucleosome that is marked with H3K4 methylation (Javasky et al. 2017). The fact that TSSs remain hyperaccessible suggests either that TSSs become occupied by nucleosomes in only a subset of cells in the population, or that nucleosomes rearrange in all cells, creating linkers between adjacent nucleosomes at TSS that are relatively large and thus more accessible than the genome-wide average. Long linkers between the nucleosomes would be expected if flanking nucleosomes reposition across the relatively large interphase nucleosome-free regions around TSSs in the absence of novo nucleosome assembly.

The level of nucleosome occupancy at the TSS is directly related to the remaining accessibility during mitosis. We find that the set of most accessible TSSs in mitosis has low levels of nucleosome occupancy, though this level is higher than in interphase. This set of most accessible TSSs also shows evidence that the TSS can frequently remain free of nucleosomes, with no changes in the positions of the directly flanking nucleosomes. One parsimonious interpretation of these data is that TSSs are variable in the extent to which nucleosomes reposition during mitosis. This variability can be at the level of single cells for individual TSSs, or at the level of subsets of TSSs where some sets remain bound by factors that maintain an open nucleosome-free site in most cells. TBP has been described as a factor that can maintain stable binding to at least a subset of promoters (Chen et al. 2002; Xing et al. 2008; Teves et al. 2018). TBP and possibly other factors may remain associated with TSSs and together with histone modifications that remain stable in mitosis serve as bookmarks for re-activation of promoters in the subsequent cell cycle. Given that we do not find any evidence that CTCF or other factors remain associated with CTCF sites in mitosis, we propose that the continued presence of modified histones and histone variants such as H2A.Z at CTCF motifs, and the larger spacing between adjacent nucleosomes around the site are sufficient for marking these sites for re-binding of CTCF as cells exit mitosis.

Both H2A.Z and H3K4 methylation marks have been studied in regard to their role as mitotic bookmarks at promoters. H2A.Z has been described to form a heterodimer with H3.3 which can fill in the NDRs of promoter regions in mitosis (Jin et al. 2009; Kelly et al. 2010; Nekrasov et al. 2012). The H3.3/H2A.Z heterodimer is found to be less stable than its canonical H3/H2A counterpart and often removed by chromosome remodelers. It has been suggested that H3.3/H2A.Z could be used as a place holder for transcription factors in mitosis. Upon mitotic exit, chromatin remodelers can be recruited to sites of H3.3/H2A.Z and open up NDRs, which will enable transcription factors to bind at its interphase sites again (Nekrasov et al. 2012). Both H3K4 monomethylation and trimethylation are maintained in mitosis ((Lin et al., 2016; Varier et al., 2010) and Fig. 5 and Supplemental Fig. S13). Recently it was observed that a large fraction of NDRs at TSSs become filled in by a nucleosome that is marked with H3K4 methylation and loss of histone acetylation (Javasky et al., 2017). Therefore, the nucleosomes that fill in NDRs stand out compared to their flanking nucleosomes that will still have acetylation marks. This could be another mechanism for specific recruitment of chromatin remodelers upon mitotic exit to these bookmarked sites, which could enable reestablishment of interphase factor binding.

We find that chromatin organization around CTCF sites alternates between two distinct states during the cell cycle. To convert from one state to the other, a number of molecular events must likely take place. First, as cells enter mitosis, CTCF dissociates from the chromatin. This likely involves phosphorylation of CTCF (Dephoure et al. 2008; Dovat et al. 2002). CTCF phosphorylation reduces its affinity for DNA in vitro (Sekiya et al. 2017; Jantz and Berg 2004). The mechanism by which nucleosomes become repositioned is not known. It is possible that simply the removal of CTCF, and associated factors such as cohesin, is sufficient for nucleosomes to passively reposition and occupy the CTCF motif. Alternatively, and more interestingly, it is possible that specific remodeling enzymes act at CTCF sites during mitosis. Sometime during or after mitotic exit CTCF regains site specific binding. This likely involves dephosphorylation of CTCF. Rebinding at CTCF motifs could be facilitated by the relative large linkers between nucleosomes around previously bound CTCF sites and the presence of histone variants and histone modifications. CTCF rebinding correlates with repositioning of the flanking nucleosomes. How CTCF rebinding and nucleosome repositioning are mechanistically linked is remains unknown. CTCF binding could passively lead to nucleosome repositioning or a process of active chromatin remodeling precedes CTCF binding.

The molecular events occurring at CTCF sites during the cell cycle coincide with large scale changes in higher order chromosomal folding, where TADs and loops are present in interphase and absent in prometaphase. The recently proposed model of loop extrusion explains how such localized events of CTCF binding can determine the formation of TADs and loops at the scale of hundreds of kilobases (Dekker and Mirny 2016; Fudenberg et al. 2018; Hansen et al. 2017). This model proposes that during interphase dynamic loop extrusion by cohesin is blocked by CTCF bound sites. This process leads to TAD formation and loops between convergent CTCF sites (Vietri Rudan et al. 2015; Wit et al. 2015; Rao et al. 2014; Guo et al. 2015). Taken together our data on CTCF and published data on cohesin (Liang et al. 2015; Waizenegger et al. 2000; Losada et al. 2002; Nagasaka et al. 2016), show that the absence of TADs and CTCF loops in mitosis can be explained by the dissociation of the entire interphase loop extrusion machinery. Interestingly, this machinery is replaced by a condensin-based loop extrusion machinery (Gibcus et al. 2018). Upon mitotic exit condensins become inactivated and CTCF and cohesin re-associate rapidly (Cai et al. 2018), allowing the reestablishment of CTCF loops and TADs.

## Methods

### Cell culturing and synchronization of culture in prometaphase

HeLa S3 cells were grown in a humidified incubator at 37C in Gibco glutamax DMEM (ThermoFisher #10566024) supplemented with 10% heat inactivated FBS (ThermoFisher # 10437028) and 5 mL Penicillin-Streptomycin (ThermoFisher #15140122) and were passaged every 2-3 days. To obtain mitotic cells, the culture at density 26,000 cells/cm^2^ was synchronized using filter sterilized 2 mM Thymidine (Sigma-Aldrich #T9250) for 24 hr, followed by a 3hr release in fresh media and a 12hr 100ng/mL nocodazole arrest (filter sterile 5000x stock in DMSO; Sigma-Aldrich #M1404). To enrich for mitotic cells, cells were gently shaking off and only rounded floating cells were collected.

HFF hTERT immortalized cells were cultured in the same media conditions as HeLa S3 cells. HFF cells were synchronized by culturing at a density of 10,000 cells/cm^2^ and arresting by a double 2mM Thymidine block of 18 and 17 hours with release in fresh media of 8 and 7 hour respectively. Cells were then synchronized in prometaphase using 100ng/mL nocodazole for 5 hours. Cells were shaken off followed by a gentle wash with dPBS. This results typically in a yield of 500-1000 mitotic cells per cm^2^ cultured cells.

We previously generated a U2OS cell line where CTCF has been homozygously and endogenously N-terminally tagged with HaloTag (FLAG-Halo-hCTCF; C32) using CRISPR/Cas9-mediated genome editing (Hansen et al., 2017a). To additionally enable the visualization of mitotic cells, here we stably integrated a previously described transgene expressing histone H2B-GFP with Puro selection (Teves et al., 2016). We transfected U2OS C32 Halo-hCTCF cells with the H2B-GFP; PURO plasmid using Lipofectamine 3000 (ThermoFisher L3000015) according to manufacturer’s protocol and then selected cells with puromycin for 1 week after which all cells in the population stably expressed H2B-GFP. U2OS cell lines stably expressing H2B-Halo-SNAP and Halo-3xNLS have been described previously (Hansen et al. 2017, 2018). The parent wild-type and C32 FLAG-Halo-hCTCF U2OS cells lines have been authenticated using STR profiling and verified to be pathogen-free (Hansen et al., 2017a).

Human U2OS osteosarcoma cells (Research Resource Identifier: RRID:CVCL_0042) were grown in a Sanyo copper alloy IncuSafe humidified incubator (MCO-18AIC(UV)) at 37C/5.5% CO2 in low glucose DMEM with 10% FBS (full recipe: 500 mL DMEM (ThermoFisher #10567014), 50 mL fetal bovine serum (HyClone FBS SH30910.03 lot #AXJ47554) and 5 mL Penicillin-streptomycin (ThermoFisher #15140122)) and were passaged every 2-4 days before reaching confluency. For live-cell imaging, the medium was identical except DMEM without phenol red was used (ThermoFisher #31053028). For live cell single-molecule imaging experiments, cells were grown overnight on 25 mm circular no 1.5H cover glasses (Marienfeld High-Precision 0117650). Prior to all experiments, the cover glasses were plasma-cleaned and then stored in isopropanol until use. For live-cell FRAP experiment and long-term time-lapse, cell preparation was identical except cells where grown on glass-bottom (thickness #1.5) 35 mm dishes (MatTek P35G-1.5-14-C).

Mitotic cells were identified through the histone H2B-GFP channel. For nocodazole prometaphase-arrested conditions, cells were treated with 100 ng/mL nocodazole (1000x stock in DMSO; Sigma-Aldrich, M1404) for 4-6 hours for imaging experiments. For “prometaphase-enriched” cells, there was no drug treatment. Instead mitotic cells with a characteristic prometaphase H2B-GFP signal were manually identified and selected for imaging. Mitotic cells for genomics studies were synchronized at a density of 16,500 cells/cm^2^ using 2mM Thymidine for 24 hrs, followed by a 3hr release in fresh media and 12hr 100ng/mL nocodazole arrest. Mitotic cells were collected by a gentle shake off.

### Flow cytometry and DAPI staining

Level of mitotic synchrony for all cultures grown for genomic studies was observed by performing flow cytometry for cell cycle analysis using propidium iodide staining of ethanol fixed. In addition to this, cells were also stained using ProLong Gold Antifade Mountant with DAPI (ThermoFisher #P36931) to determine the fraction of prometaphase cells in total culture based on chromatin condensation.

### Mitotic chromatin cluster purification

Mitotic chromatin clusters were purified according to previously a published protocol with minor changes (Gasser and Laemmli 1987). Cells were synchronized in prometaphase using thymidine and nocodazole arrests as described above. After washing cells in dPBS, cells were lysed using 0.1% NP-40, 10% glycerol, 15 mM Tris-HCl, 80 mM KCl, 2mM EDTA-KOH, 0.2 mM spermine, 0.5 mM spermidine, 1mM DTT and Roche cOmplete EDTA-free protease inhibitor (Sigma-Aldrich 11873580001). Lysis was followed by dounce homogenization using pestle B. After proper lysis and removal of cytosolic fractions was obtained, the solution was transferred to percoll gradients, containing a bottom layer of 60% percoll plus (Sigma-Aldrich #E0414), 15% Glycerol, 1 mM DTT, Roche cOmplet EDTE-free protease inhibitor, 0.1% NP-40, 15 mM Tris-HCl, 80 mM KCl, 2mM EDTA-KOH, 0.8 mM Spermidine and 2 mM Spermine, and a top layer of 25% Glycerol, 0.1% NP-40, 15 mM Tris-HCl, 80 mM KCl, 2mM EDTA-KOH, 1 mM DTT, Roche cOmplet EDTE-free protease inhibitor, 0.1 mM Spermidine and 0.25 mM Spermine. Gradients were spun at 1000xg for 30 minutes to separate mitotic chromatin and interphase nuclei. Mitotic chromatin pellets on top of the bottom percoll layer. The top layer was taken off to remove interphase nuclei. Pestle B was again used to dounce mitotic chromatin clusters and the solution was diluted 1 to 4 with 60% percoll plus, 15% Glycerol, 1 mM DTT, Roche cOmplete EDTA-free protease inhibitor, 0.1% NP-40, 15 mM Tris-HCl, 80 mM KCl, 2mM EDTA-KOH, 0.8 mM Spermidine and 2 mM Spermine. The solution was spun down at 3000xg for 5 minutes and then at 44,500xg for 30 minutes. The supernatant was again removed up till 2-3 cm and pelleted chromatin clusters were diluted in 15 mM Tris-HCl, 80 mM KCl, 2mM EDTA-KOH, 1mM DTT, 0.1% NP-40, 0.05 mM spermine, 0.125 mM spermidine and Roche cOmplete EDTA-free protease inhibitor. Mitotic chromatin clusters were washed several times with this solution to remove remaining percoll plus. Mitotic chromatin clusters can be frozen at −80C in 33.33% glycerol. Chromatin clusters were used for ATAC-seq both directly after chromatin cluster purification and after freezing. Supplemental Fig. S14 shows fresh or frozen clusters obtain highly similar results for ATAC-seq. For this reason we continued using frozen chromatin clusters for all further ATAC-seq experiments. Mitotic chromatin clusters for 5C were immediately fixed in 1% formaldehyde before pelleting and storage at −80C.

### 5C

For each 5C library, 25 million cells or the equivalent in mitotic chromatin clusters from HeLa S3 were used as input. There were two biological replicates produced for each condition. Cells were crosslinked for 10 minutes at room temperature in a final concentration of 1% formaldehyde. Glycine was added to quench crosslinking and cells were incubated for 5 min with gentle agitation and then on ice for 15 min. Cells were spun down and after removal of supernatant, pellets were frozen at −80C and stored until used for 5C. 5C was performed according to published protocols (Hnisz et al. 2016; Dostie et al. 2006). We investigated the same regions as previously described and we used the same primer pool (Hnisz et al. 2016). This 5C primer pool covers two 2 Mb regions located on chromosome 1 (hg19 chr1: 46740122-48740121) and chromosome 11 (hg19 chr11: 33003550-35003549). Cells were lysed in presence of protease inhibitors and a dounce homogenizer with pestle A was used to obtain proper removal of the cytosolic fraction. Purified mitotic chromatin clusters did not need to be lysed, and were simply resuspended and washed in 1xNEBuffer 2.1 (New England Biolabs #B7202). Chromatin for all conditions was digested using 400U HindIII (New England Biolabs #R0104) and incubated at 37C overnight. After digestion the enzyme was inactivated, followed by ligation for 2 hours at 16C with 10U of T4 ligase (Invitrogen 15224017) in presence of 1% Triton X-100. Proteins were digested using proteinase K, followed by phenol chloroform extraction, ethanol precipitation and RNA digestion using RNase A to purify DNA and obtain 3C templates to continue the 5C protocol. Different conditions were tested to optimize genome copy number, 5C probe concentration and number of amplification cycles. From this, we decided to use ~800,000 genome copies, 0.3 fmole probes and 20 cycles of PCR amplification. Annealing of the 5C probes was performed at 50C overnight, directly followed by ligation of the probes at 50C for 1 hour using Taq ligase (New England Biolabs #M0208). After primer annealing and ligation, 5C ligation products were amplified for 20 cycles and libraries were size selected by gel purification to remove primer dimers and unbound primers. The 5C libraries were then prepared for sequencing using barcoded adaptors and TruSeq Nano DNA sample kit (Illumina 20015964), followed by final amplification with 6 cycles. The libraries were again gel purified and finally sequenced on the Illumina HiSeq4000 platform to obtain 50 bp paired-end reads.

Sequencing data was processed using a custom pipeline for mapping and assembly of 5C interactions, as previously described (Lajoie et al. 2009). After careful comparison, data from two biological replicates were combined to produce a single interaction map. Mapping statistics for each library can be found in table S1. 5C interaction matrices were filtered to remove the diagonal, flagged outlier singletons and rows/columns based on Z-score cut-off of 12 for both cis and trans interactions. When singletons or row/columns had to be removed from an individual 5C matrix, we removed this singleton or row/column from every of our 5C matrices. As the primers used for this study are designed using the double-alternating pattern, we merged rows/columns that map to the same restriction fragment using the mean of the interactions and the interaction matrices were made symmetrical. Data was balanced using ICE, binned in 20 kb bins and balanced again (Imakaev et al. 2012). Insulation index and 5C anchor plots were calculated using published methods (Crane et al. 2015; Sanyal et al. 2012). Data was balanced and binned at 15kb for 5C anchor plot analysis.

### ATAC-seq

ATAC-seq was performed following a previously published protocol (Buenrostro et al. 2015). Briefly, 50,000 cells per experiment (typically 2 biological replicates, 4 technical replicates per dataset) were washed and lysed using lysis buffer containing 0.1% NP-40, 10 mM Tris-HCl (pH 7.4), 10 mM NaCl and 3 mM MgCl2.. Cells were transposed using the Nextera DNA library prep kit (Illumina #FC-121-1030) for 30 min at 37C. DNA was immediately purified using Qiagen MinElute Kit (Qiagen #28004). Cycle number for PCR amplification of the DNA was determined for each sample individually using qPCR. Primers were removed from amplified libraries using AMpure XP beads (Beckman Coulter #A63881). The libraries were sequenced on HiSeq2000 or HiSeq4000 illumina sequencers to obtain 2×50 paired-end reads.

ATAC-seq data were trimmed from their 3’ end to 24 bp and aligned to hg19 using bowtie2 with maximum mapping length of 2000 bp (Langmead and Salzberg 2012). Paired-end reads that mapped with a mapping quality below 3 were discarded, as were reads mapped to the mitochondrial genome and PCR duplicates. In addition, regions marked as DAC blacklisted regions in hg19 by the ENCODE project were removed (annotation file accession number ENCSR636HFF). Reproducibility between technical replicates was evaluated based on similarities in length distribution, percentage non-valid reads (e.g. percentage mitochondrial DNA, PCR duplicates etc), genome track and peak calling before combining data. For further analysis (unless otherwise indicated) each end +5 bp was taken and analysis was continued treating the reads as single end reads. Bedtools was used to produce bigWig tracks for several bin sizes, as well as for manipulation of other bed files (Quinlan and Hall 2010). Peak calling was performed using HOMER (Heinz et al. 2010). To group TSSs in top 20% highest accessible sites and top 20% lowest accessible sites, all peaks in mitotic chromatin at TSSs were ranked based on HOMER-score (reflecting signal over background enrichment).

In order to produce V-plots, paired-end reads for each library were separated based on total read length into bins of 25 bp. For each bin, HOMER aggregation plots for an element of interest (e.g. CTCF motifs with interphase ATAC-seq peaks) were produced using only the midpoint of the read. The values of the aggregation plots were used to produce heatmaps. Each read length bin was normalized using HOMER normalization for genome wide coverage in order to visualize the rare larger length bins, ensuring the sum of the reads in each length bin genome wide is comparable. V-plots of unnormalized length bins and V-plots of a randomized region are shown in Supplemental Fig. S15. For CTCF-centered V-plots, all CTCF motifs were aligned in the same direction. For TSS-centered V-plots, all TSS were in the same direction.

### CTCF motif identification for hg19

A candidate list of all CTCF motifs in the human genome was produced using CTCF motif consensus for vertebrates published in 2007 by Kim et al. as an input for HOMER (Kim et al. 2007; Heinz et al. 2010). Orientation of CTCF motif was always taken into account during analysis.

### ENCODE data sets

ENCODE ChIP-seq data for CTCF in HeLa S3 was downloaded for analysis. GEO accession #SRX067536.

### CUT&RUN

CUT&RUN was performed according to the published protocol with minor changes (Skene and Henikoff 2017). As mitotic cells do not have a nuclear membrane, it was not possible to use concanavalin A beads. Instead, cells were spun at 600xg for 3 min at 4C for every wash or buffer exchange. Asynchronous and mitotic cells were harvested fresh for every CUT&RUN experiment. 500,000 cells were used as input for every time point and there were 2 biological replicates produced for every dataset. Cells were lysed on ice for 10 minutes in 20mM HEPES-KOH pH 7.9, 10mM KCl, 0.5mM spermidine, 0.1% Triton-X-100, 20% glycerol and Roche cOmplete EDTA-free protease inhibitor (Sigma-Aldrich 11873580001). Cells were washed in wash buffer 1 (20mM HEPES pH 7.5, 150mM NaCl, 2mM EDTA 0.5mM spermidine, 0.1% BSA and Roche cOmplete EDTA-free protease inhibitor) and then resuspended in wash buffer 2 (20mM HEPES pH 7.5, 150mM NaCl, 0.5mM spermidine, 0.1% BSA and Roche cOmplete EDTA-free protease inhibitor). Cells were incubated with primary antibody according to table S2 for 2 hours while rotating at 4C. A control, for which no primary antibody was used at this step, was always included in each experiment. Cells were washed several times using wash buffer 2, followed by incubation with 1:400 pA-MNase (batch #6 from Henikoff lab) for 1 hour on rotator at 4C. Cells were washed several times in wash buffer II, after which cell suspensions were split in 150 μL wash buffer II per time-point sample. Samples were transferred to an ice water bath equilibrated at 0 degrees Celsius. MNase digestion was activated by adding CaCl_2_ to final concentration of 2mM while vortexing. MNase was inactivated after designated time (typically 5 seconds, and up to 30 minutes) by adding 2XSTOP+ (200mM NaCl, 20mM EDTA, 4mM EGTA, 50μg/mL RNaseA and 40μg/mL Glycogen). The samples were then incubated for 20 minutes at 37C to digest RNA, followed by adding 0.1% SDS and proteinase K and 10 minutes incubation at 65C. DNA was extracted from the samples using 1:1 phenol chloroform extraction, followed by size selection to remove large fragments using AMpure XP beads (Beckman Coulter #A63881). DNA was purified using ethanol precipitation and samples were finally dissolved in Tris-EDTA. Samples were prepared for sequencing using barcoded adaptors of TruSeq Nano DNA Sample Prep Kit (Illumina 20015964) and amplified. Finally libraries were size selected using gel purification for an insert size of 50-200 bp. DNA was purified from gel using QIAquick gel extraction kit (Qiagen #28704). CUT&RUN libraries were sequenced on Illumina MiSeq sequencer to obtain 50 bp paired-end reads to a sequence depth of 1 to 5 million valid reads per sample.

CUT&RUN sequenced reads were trimmed from 3’ end to 24 bp and aligned to hg19 using bowtie2 with maximum mapping length of 2000 bp (Langmead and Salzberg 2012). Paired-end reads that mapped with a mapping quality below 3 were discarded, as well as PCR duplicates. Paired-end reads were separated based on total read length above or below 120 bp. HOMER was used to produce aggregation plots on elements of interest and to perform de novo peak calling on CUT&RUN data (Heinz et al. 2010).

### Western blot

Cells and mitotic clusters were harvested, washed in dPBS and pelleted followed by freezing and storage at −80C until used. Protein lysates were obtained using RIPA buffer (ThermoFisher #89900) supplemented with 1x HALT protease inhibitor (ThermoFisher #78429). All samples were incubated on ice for 15 minutes while firmly pipetting every couple of minutes. Samples were spun down 30 min at 16,100xg at 4C and supernatant was collected. Pierce Lane Marker Reducing Sample buffer (ThermoFisher #39000) was added and protein lysate was incubated for 30 min at 95C. Samples to detect CTCF were ran on a 3-8% Tris-Acetate gel, whereas samples for histone detections were ran on a 4-12% Tris-Bis gel. Antibodies used in this study as well as their concentration used for western blot are listed in table S2. Primary antibody incubation was done overnight, followed by secondary antibody incubation by either goat anti-rabbit IgG-HRP (Cell Signaling Technologies #7074) or goat anti-mouse IgG-HRP (Cell Signaling Technologies #7076). HRP was activated using SuperSignal West Dura Chemiluminescent Substrate (ThermoFisher #34076) and signal was detected using the BioRad Gel Doc system.

### Fluorescence Recovery After Photobleaching (FRAP) imaging

FRAP experiments were performed and analyzed as previously described (Hansen et al. 2017). Briefly, FRAP was performed on an inverted Zeiss LSM 710 AxioObserver confocal microscope equipped with a motorized stage, a full incubation chamber maintaining 37°C/5% CO_2_, a heated stage, an X-Cite 120 illumination source as well as several laser lines. Halo-TMR was excited using a 561 nm laser and H2B-GFP was excited using a 488 nm laser. Images were acquired on a 40x Plan NeoFluar NA1.3 oil-immersion objective at a zoom corresponding to a 100 nm x 100 nm pixel size and the microscope controlled using the Zeiss Zen software. In FRAP experiments, 300 frames were acquired at either 1 frame per second allowing 20 frames to be acquired before the bleach pulse to accurately estimate baseline fluorescence. A circular bleach spot (r = 10 pixels) was chosen in a region of homogenous fluorescence at a position at least 1 μm from nuclear boundaries and in a region occupied by chromatin as visualized using the H2B-GFP transgene in the GFP channel. The spot was bleached using maximal 561 nm laser intensity and pixel dwell time corresponding to a total bleach time of ~1 s. We generally collected data from several cells per cell line per condition per day and all presented data is from at least three independent replicates on different days: U2OS C32 Halo-hCTCF interphase: 32 cells; prometaphase-arrested (100 ng/mL nocodazole for 4-6 hours) U2OS C32 Halo-hCTCF, H2B-GFP (19 cells); prometaphase-enriched (no drug treatment; prometaphase-like cells identified through the histone H2B-GFP channel) U2OS C32 Halo-hCTCF, H2B-GFP (18 cells) U2OS Halo-3xNLS in interphase (6 cells); and U2OS H2B-Halo-SNAP prometaphase-enriched (no drug treatment; prometaphase-like cells identified through the histone H2B-TMR channel; 10 cells). To quantify and drift-correct the FRAP movies, we used a previously described custom-written pipeline in MATLAB (Hansen et al. 2017). Briefly, we manually identify the bleach spot. The nucleus is automatically identified by thresholding images after Gaussian smoothing and hole-filling (to avoid the bleach spot as being identified as not belonging to the nucleus). We use an exponentially decaying (from 100% to ~85% (measured) of initial over one movie) threshold to account for whole-nucleus photobleaching during the time-lapse acquisition. Next, we quantify the bleach spot signal as the mean intensity of a slightly smaller circle (*r* = 0.6 μm), which is more robust to lateral drift. The FRAP signal is corrected for photobleaching using the measured reduction in total nuclear fluorescence (~15% over 300 frames at the low laser intensity used after bleaching) and internally normalized to its mean value during the 20 frames before bleaching. We correct for drift by manually updating a drift vector quantifying cell movement during the experiment. Finally, drift- and photobleaching corrected FRAP curves from each single cell were averaged to generate a mean FRAP recovery. We used the mean FRAP recovery in all figures and error bars show the standard error of the mean.

### Single-molecule imaging (spaSPT)

U2OS C32 FLAG-Halo-hCTCF; H2B-GFP cells grown overnight on plasma-cleaned 25 mm circular no 1.5H cover glasses (Marienfeld High-Precision 0117650). After overnight growth, cells were either treated with nocodazole or not and then labeled with 50 nM PA-JF_549_ (Grimm et al. 2016) for ~15-30 min and washed twice (one wash: medium removed; PBS wash; replenished with fresh medium). At the end of the final wash, the medium was changed to phenol red-free medium keeping all other aspects of the medium the same (and adding back 100 ng/mL nocodazole if appropriate). Single-molecule imaging was performed on a custom-built Nikon TI microscope equipped with a 100x/NA 1.49 oil-immersion TIRF objective (Nikon apochromat CFI Apo TIRF 100x Oil), EM-CCD camera (Andor iXon Ultra 897; frame-transfer mode; vertical shift speed: 0.9 μs; −70°C), a perfect focusing system to correct for axial drift and motorized laser illumination (Ti-TIRF, Nikon), which allows an incident angle adjustment to achieve highly inclined and laminated optical sheet illumination (Tokunaga et al. 2008). An incubation chamber maintained a humidified 37°C atmosphere with 5% CO_2_ and the objective was also heated to 37°C. Excitation was achieved using a 561 nm (1 W, Genesis Coherent) laser for PA-JF_549_. Histone H2B-GFP was visualized using epi-illumination (X-cite 120LED). The excitation laser was modulated by an acousto-optic tunable filter (AA Opto-Electronic, AOTFnC-VIS-TN) and triggered with the camera TTL exposure output signal. The laser light was coupled into the microscope by an optical fiber and then reflected using a multi-band dichroic (405 nm/488 nm/561 nm/633 nm quad-band, Semrock) and then focused in the back focal plane of the objective. Fluorescence emission light was filtered using a single band-pass filter placed in front of the camera using the following filters (PA-JF549: Semrock 593/40 nm bandpass filter). The microscope, cameras, and hardware were controlled through NIS-Elements software (Nikon).

We recorded single-molecule tracking movies using our previously developed technique, stroboscopic photo-activation Single-Particle Tracking (spaSPT) (Hansen et al. 2018, 2017). Briefly, 1 ms 561 nm excitation (100% AOTF) of PA-JF_549_ was delivered at the beginning of the frame; 405 nm photo-activation pulses were delivered during the camera integration time (~447 μs) to minimize background and their intensity optimized to achieve a mean density of ~1 molecule per frame per nucleus. 30,000 frames were recorded per cell per experiment. The camera exposure time was 7 ms resulting in a frame rate of approximately 134 Hz (7 ms + ~447 μs per frame).

spaSPT data was analyzed (localization and tracking) and converted into trajectories using a custom-written Matlab implementation of the MTT-algorithm (Sergé et al. 2008) and the following settings: Localization error: 10^−6.25^; deflation loops: 0; Blinking (frames): 1; max competitors: 3; max *D* (μm^2^/s): 20.

We considered three conditions and recorded ~7-8 cells per conditions per replicate and performed three independent replicates on three different days. Specifically, the conditions were U2OS C32 Halo-hCTCF; H2B-GFP in interphase (22 cells total); prometaphase-arrested (100 ng/mL nocodazole for 4-6 hours) U2OS C32 Halo-hCTCF; H2B-GFP (24 cells); and prometaphase-enriched (no drug treatment; prometaphase-like cells identified through the histone H2B-GFP channel) U2OS C32 Halo-hCTCF; H2B-GFP (23 cells).

### Model-based analysis of single-molecule tracking data using Spot-On

To analyze the spaSPT data, we used our previously described kinetic modeling approach (Spot-On) (Hansen et al. 2018, 2017). Briefly, we analyze each replicate separately and the reported bound fractions and free diffusion coefficients are reported as the mean +/− standard deviation from analyzing each replicate separately. We merged the data from all cells (~7-8) for each replicate, compiled histograms of displacements and then fit the displacement cumulative distribution functions for 8 timepoints using a 2-state model that assumes that CTCF can either exist in a chromatin-bound or freely diffusive state:

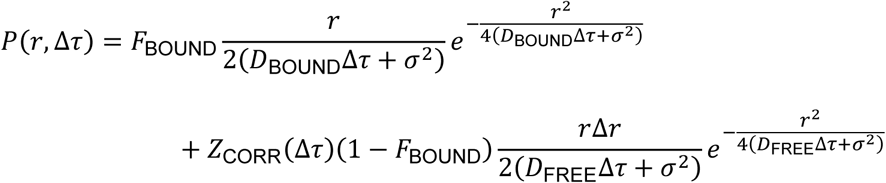
where:

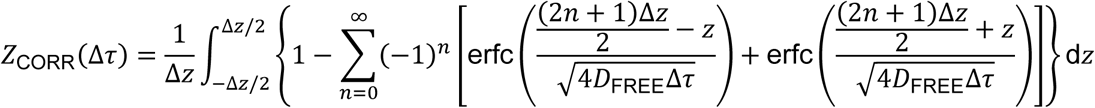
and:

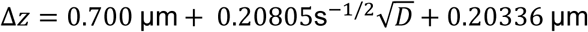

Here, *F*_BOUND_ is the fraction of molecules that are bound to chromatin, *D*_BOUND_ is diffusion coefficient of chromatin bound molecules, *D*_FREE_ is diffusion coefficient of freely diffusing molecules, *r* is the displacement length, Δ*τ* is lag time between frames, *Δz* is axial detection range, *σ* is localization error and *Z*_CORR_ corrects for defocalization bias (i.e. the fact that freely diffusion molecules gradually move out-of-focus, but chromatin bound molecules do not). Model fitting and parameter optimization was performed using a non-linear least squares algorithm (Levenberg-Marquardt) implemented in the Matlab version of Spot-On (v1.0; GitLab tag 92cdf210) and the following parameters: dZ=0.7 μm; GapsAllowed=1; TimePoints: 8; JumpsToConsider=4; ModelFit=2; NumberOfStates=2; FitLocError=1; D_Free_2State=[0.4;25]; D_Bound_2State=[0.00001;0.05];

### Spot-On data and computer code availability

The raw spaSPT data is available in Spot-On readable Matlab and CSV formats in the form of single-molecule trajectories at https://zenodo.org/record/1306976. The Spot-On Matlab code is available together with a step-by-step guide at Gitlab: https://gitlab.com/tjian-darzacq-lab/spot-on-matlab. For additional documentation see also the Spot-On website https://SpotOn.berkeley.edu and previous publications (Hansen et al. 2018, 2017).

### Time-lapse imaging

Multi-color long-term time-lapse imaging of U2OS C32 Halo-hCTCF; H2B-GFP was performed using a Nikon BioStation motorized inverted microscope equipped with a 40x 0.8 NA Plan Fluor objective and filter sets appropriate GFP and TMR and a DS-Qi1 CCD-camera. The BioStation maintains incubation conditions (37⎕C/5% CO2) and remains mechanically stable for hours. Time-lapse movies were recorded using phase, GFP (for H2B-GFP) and TMR (for TMRHalo-hCTCF) acquiring one frame every 2 or 5 minutes and lasted for a total time of at least 10 hours – enough that most cells in a field of view went through mitosis. CTCF was clearly enriched on mitotic chromosomes during most phases of mitosis.

### Data access

5C, ATAC-seq and CUT&RUN data will be publically available at GEO upon publication (GEO accession number in process). All imaging data is publically available at Zenodo (https://zenodo.org/record/1306976).

## Acknowledgments

We especially thank Sarah Hainer and Tom Fazzio for advice on ATAC-seq data interpretation and providing reagents. We also thank Bas van Steensel for advice and suggestions. We thank Sheila Teves and all current and former Dekker lab members for helpful discussions; especially Betul Akgol Oksuz, Bryan Lajoie, Ankita Nand and Hakan Ozadam for advice on data analysis and Allana Schooley for helpful comments regarding the manuscript. This work was supported by grants from the National Human Genome Research Institute (HG003143) and the NIH Common Fund (DK107980) to J.D. and by NIH grants UO1-EB021236 and U54-DK107980 (X.D.), the California Institute of Regenerative Medicine grant LA1-08013 (X.D.). This work was performed in part at the UC Berkeley CRL Molecular Imaging Center, supported by the Gordon and Betty Moore Foundation. A.S.H. is a postdoctoral fellow of the Siebel Stem Cell Institute. J.D. is an investigator of the Howard Hughes Medical Institute.

## Author contributions

M.E.O. and J.D. conceived and designed the project. Y.L and M.E.O. preformed 5C. M.E.O. generated ATAC-seq and CUT&RUN datasets. A.S.H. generated U2OS cell line and obtained and analyzed all imaging data. M.E.O. analyzed 5C, ATAC-seq and CUT&RUN data-sets. M.E.O., A.S.H., X.D. and J.D. wrote the manuscript.

